# CRISPR targeting of FOXL2 c.402C>G mutation reduces malignant phenotype in granulosa tumor cells and identifies anti-tumoral compounds

**DOI:** 10.1101/2024.07.01.601520

**Authors:** Sandra Amarilla-Quintana, Paloma Navarro, Alejandra Ramos, Ana Montero-Calle, Pablo Cabezas-Sainz, Maria J Barrero, Diego Megías, Borja Vilaplana-Martí, Carolina Epifano, Déborah Gómez-Domínguez, Iván Hernández, Sara Monzón, Isabel Cuesta, Laura Sánchez, Rodrigo Barderas, Jesús García-Donas, Alberto Martín, Ignacio Pérez de Castro

## Abstract

FOXL2 is a transcription factor essential for sex determination and ovary development and maintenance. Mutations in this gene are implicated in syndromes involving premature ovarian failure and granulosa cell tumors (GCTs). This rare cancer accounts for less than 5% of diagnosed ovarian cancers and is causally associated with the FOXL2 c.402C>G, p.C134W mutation in 97% of the adult cases (AGCTs). In this study, we employed CRISPR technology to specifically eliminate the FOXL2 c.402C>G mutation in granulosa tumor cells. Our results show that this Cas9-mediated strategy selectively targets the mutation without affecting the wild-type allele. Granulosa cells lacking the FOXL2 c.402C>G exhibit a reduced malignant phenotype, with significant changes in cell proliferation, invasion, and cell death. Furthermore, these modified cells are more susceptible to Dasatinib and Ketoconazole. Transcriptomic and proteomic analyses reveal that CRISPR-modified granulosa tumor cells shift their expression profiles towards a wild-type like phenotype. Additionally, this altered expression signature has led to the identification of new compounds with antiproliferative and pro-apoptotic effects on granulosa tumor cells. Our findings demonstrate the potential of CRISPR technology for the specific targeting and elimination of a mutation causing granulosa cell tumors, highlighting its therapeutic promise for treating this rare ovarian cancer.

**Simple Summary:** Adult granulosa cell tumors (AGCTs), characterized by a specific point mutation (C134W) in the gene FOXL2, are less than 5% of all the diagnosed ovarian cancers. Surgery is the cornerstone treatment for AGCT even at relapse, with systemic therapy showing poor results. The aim of our study is to explore the potential therapeutic effect of the elimination of the C134W mutation. To achieve our goal, we have eliminated the mutant allele using CRISPR technology. Our results demonstrate that CRISPR-mediated elimination of FOXL2-C134W reduces the malignant phenotype of granulosa tumor cells, which change their transcriptional, proteomic, and cellular phenotype to a wild type like, granulosa type. Moreover, the induced changes allowed us to find new compounds with antitumoral activity. This work highlights the therapeutic potential of CRISPR mediated technology for the treatment of AGCT.

## Introduction

Granulosa cell tumors (GCTs) are a rare type of cancer, constituting less than 5% of all the ovarian neoplasms and representing the most common sex-cord stromal derived tumors [1,2]. GCTs are classified into juvenile (JGCTs) or adult (AGCTs) types based on clinical and histopathological characteristics. JGCTs, diagnosed primarily in patients under 30 years, are very infrequent, accounting for 5% of GCTs cases. AGCTs are more common, predominantly affecting perimenopausal women. AGCTs originate from granulosa cells, which are crucial for follicle maturation and the synthesis of inhibin, estradiol, and anti-Müllerian hormone (AMH). While these hormones are normally secreted in premenopausal women, elevated serum levels of inhibin B and AMH in AGCTs patients serve as significant tumor markers [3]. Other histopathological markers used to determine tumor type and stage include androgen receptor, calretinin, SF-1, and FOXL2 [4,5].

Despite most AGCT cases are diagnosed at stage I due to the slow growth of these tumors, advanced stages decrease the 10-year survival rate and increase relapse risk [2]. Initial treatment typically involves surgery, including total hysterectomy and bilateral salpingo-oophorectomy in older women, with more conservative surgery options for those younger women with early-stage GCT. Adjuvant chemotherapy with platinum-based combinations is frequently administered for locally advanced disease, though its benefits remain unclear. Recurrences can occur decades after initial surgery, typically involving the peritoneal cavity, though visceral metastasis have also been reported [6–8]. Platinum-based chemotherapy may induce partial responses in recurrent disease but is not curative [9]. Various drugs, including tyrosine-kinase inhibitors and hormone therapies like aromatase inhibitors, have been evaluated in clinical trials and series of cases [10–12]. Unfortunately, disease stabilization rather than significant tumor reduction is often observed, likely due to the slow growth of AGCTs rather than drug efficacy.

The most prominent molecular characteristic of AGCTs is a single-nucleotide mutation in the FOXL2 gene, present in 95-97% of the cases and first described in 2009 [13]. This somatic mutation, occurring in heterozygosity, is a missense point mutation (C402G; Cys134Trp) specific to AGCTs and rarely seen in JGCTs [14]. FOXL2, a forkhead transcription factor, is mainly expressed in granulosa cells and eyelids in mammals and is associated with BPES syndrome type I (Blepharophimosis-Ptosis-Epicanthus) [15–17] and POF (premature ovarian failure) [18]. Depletion of FOXL2 leads to a lack of primordial follicles, due to granulosa cell differentiation problems that induce atresia, affecting directly to the female fertility [19]. In animal models and during embryonic stages, FOXL2 has been described as a marker of ovarian differentiation [20,21]. In fact, FOXL2 is implicated in ovary development and maintenance through the regulation of PI3K, ERK and Smad pathways, among others [22–24]. Furthermore, FOXL2 is involved in relevant biological processes related to cholesterol metabolism, apoptosis, cell proliferation, and differentiation, as well as immunomodulation [25]. The FOXL2 c.402C>G, p.C134W mutation alters the FOXL2 DNA-binding domain, leading to alterations in its transcriptional regulatory capacity. Thus, FOXL2 C134W is described to form dimers with Smad proteins, a crucial step for inducing the transcriptional activation of genes involved in survival, proliferation and epithelial to mesenchymal transition, which culminates in the development of AGCTs [26].

Given the presence of this single-nucleotide mutation, CRIPSR-Cas9 technology is an ideal tool to investigate the impact of deleting FOXL2 c.402C>G on tumor phenotype and to explore the underlying molecular mechanisms. Understanding these aspects can lead to new therapeutic strategies, potentially improving survival and health outcomes of AGCT patients.

In this study, we demonstrate that eliminating FOXL2 c.402C>G induces anti-tumor properties, increases sensitivity to certain therapeutic agents and identifies dysregulated pathways and processes that could be targeted by repurposed drugs.

## Materials and Methods

Details of reagents, kits, plasmid, antibodies, and other resources included in this work are listed in Table 1.

**Table 1.**
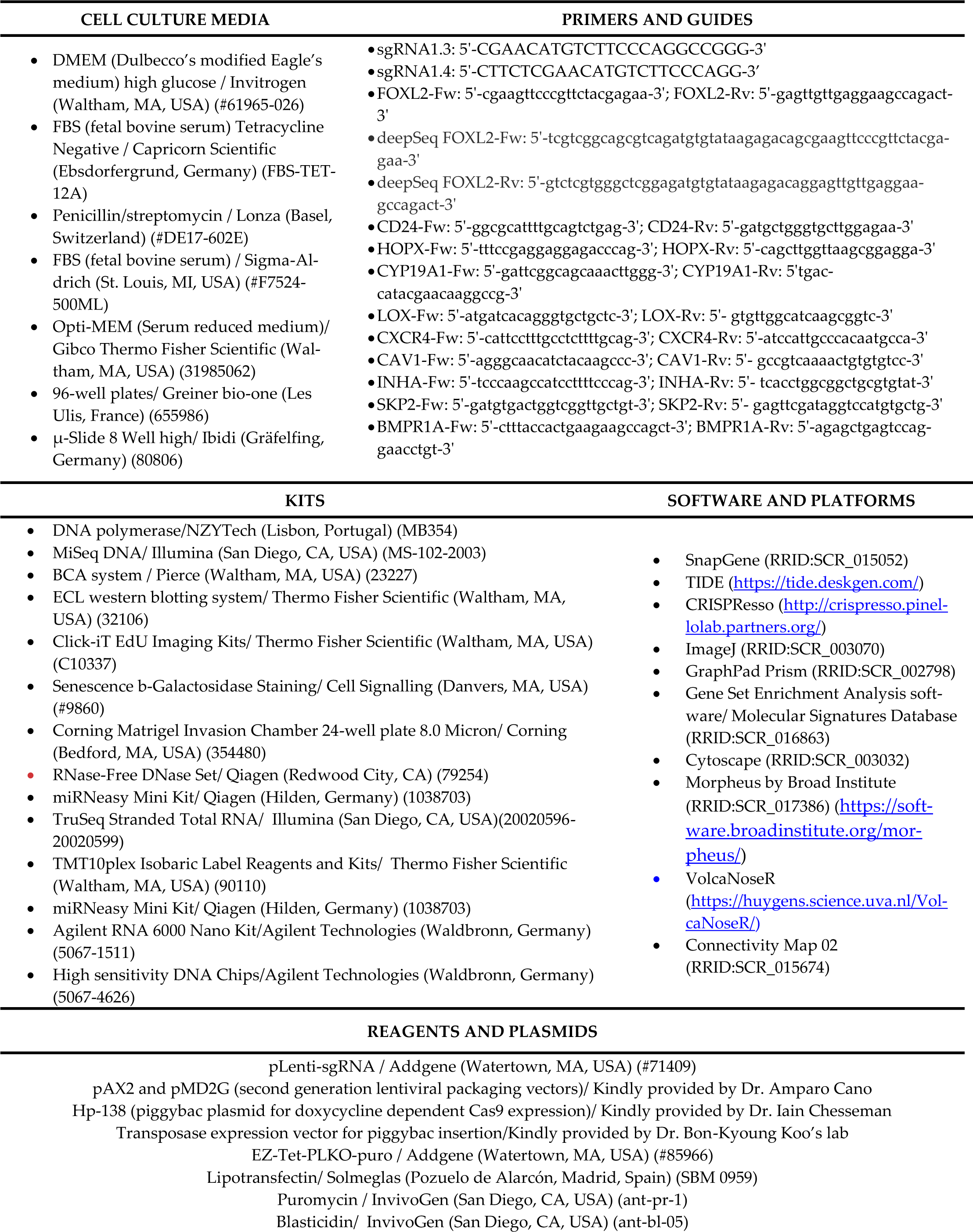
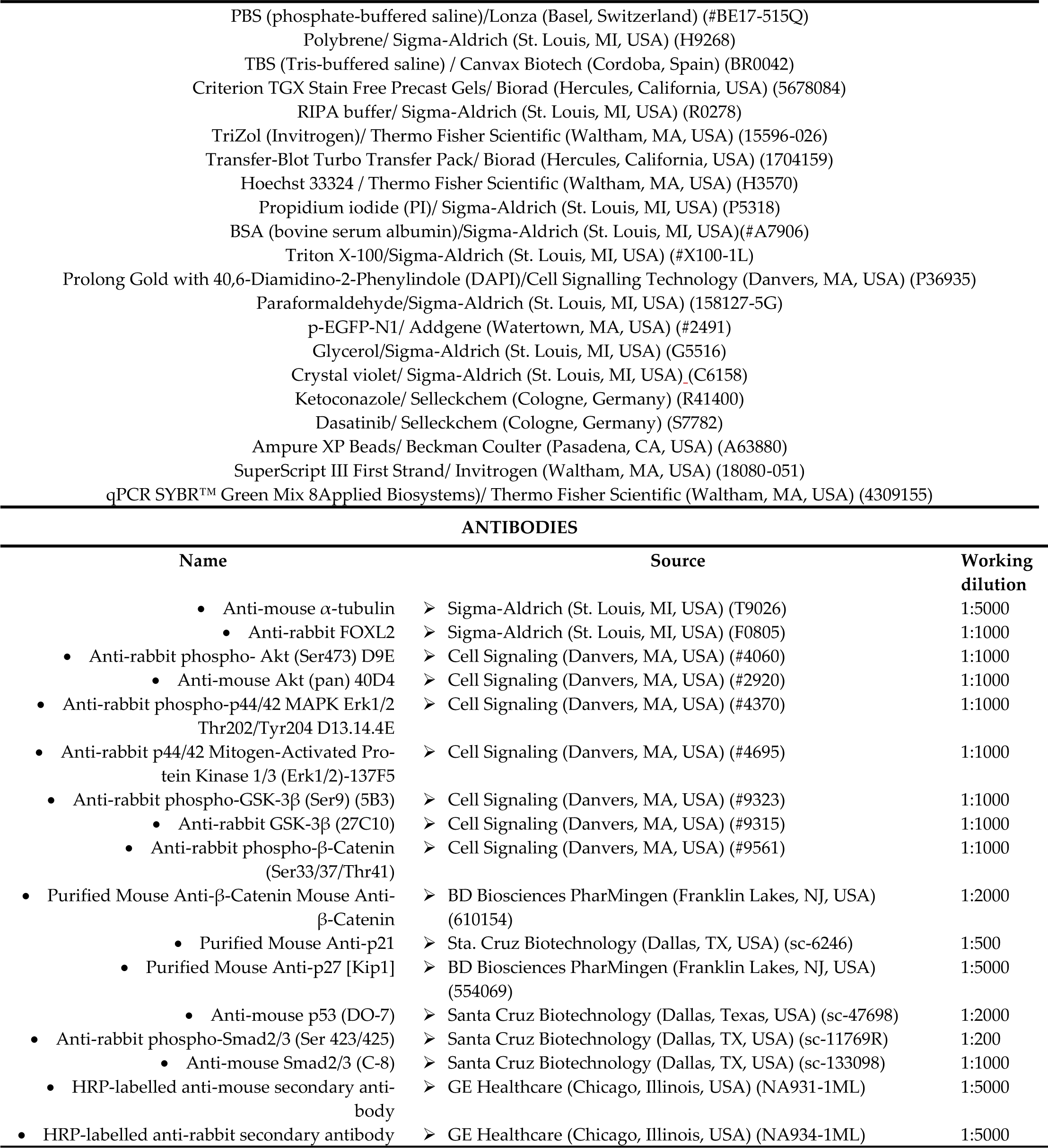
Resources used in this work.

### Cell lines

We have used the KGN cell line (Riken BioResource; RCB1154) stablished from a 63-year-old Japanese ovarian granulosa cell tumor patient [27]. Parental cell KGN and stablished clones derived from the same cell line were cultured at 37°C and 5% CO_2_ in DMEM medium (Dulbecco Modified Eagle Medium) supplemented with 10% bovine fetal serum (FBS) and 1% of penicillin/streptomycin. HEK293T cell line was used to generate lentiviral particles. Culture medium was also DMEM, supplemented with FBS 10%. Cells grew at 37°C and 5% CO_2_.

### Clone generation using CRISPR/Cas9 and genomic characterization

Single guides were designed to target the mutated allele of FOXL2, using Breaking Cas Design tool (https://bioin-fogp.cnb.csic.es/tools/breakingcas/). Guides sg1.3 and sg1.4 (Table 1) were cloned in the pLenti-sgRNA plasmid for virus generation and cell infection. Lentivirus containing spCas9 sequence or one of the two specific FOXL2 C134W mutation guides (sg1.3 or sg1.4) were generated to infect KGN cell line. HEK293T cells were seeded 24 hours before transfection in medium lacking penicillin/streptomycin. For transfection, either plasmid containing spCas9, or one of the two guides together with the second-generation packaging vectors, were mixed with lipotransfectin and added to the HEK293T cells medium. Viruses from the different conditions were collected after 48and 72-hours post-transfection, centrifuged at 1200 rpm for 10 minutes and viral supernatant was filtered with a 0,45 µm filter and stored at -80 ͦ. For infection, supernatant containing spCas9 lentivirus was mixed with 8 µg/ml polybrene, added to the KGN culture and incubated at 37 ͦ C and 5% CO2 for at least 6 hours. After that time, fresh DMEM medium was added to the culture. Selection of cells containing Cas9 plasmid required the presence of 5 µg/ml of blasticidin in the medium for 13 days. The same infection protocol was used to infect KGN cells with lentiviral particles harboring sg1.3 or sg1.4 guides. In this case, 48 hours post infection, selection was done with 2 µg/ml of puromycin for 48 hours. After completing antibiotic treatment, 3 different pools were generated (no guide, sg1.3 and sg1.4) and subsequently divided in two. One part of the culture was seeded, by serial dilution, in p96 well plates containing 1 cell per well to generate individual clones. The remaining part was kept as bulk culture (named as “pool”), which was used to study the activity and the specificity of the guides.

To determine CRISPR/Cas9 activity, DNA from pools and clones was isolated. Amplification of the sequence including the mutation site was performed by PCR using FOXL2-Fw and FOXL2-Rv primers and DNA polymerase. Sanger sequencing was carried out in the Genomic Unit of Instituto de Salud Carlos III (ISCIII) and then combined with the TIDE platform to determine the presence and magnitude of insertions and deletions present in the analyzed PCR products. To specifically determine the novel allelic variants, MiSeq DNA was performed on the target region PCR amplified with primers deepSeq FOXL2-Fw and deepSeq FOXL2-Rv (Table 1). The resultant PCR products were used for a second PCR to add specific indexes required for ulterior sample identification. FASTQ files comprising sequencing data were analyzed by CRISPResso. This platform analyzes the region closest to the Cas9 cut and compares the different insertions/deletions/substitutions found in the analyzed sample with a reference sequence.

A nucleofection strategy was also used to deliver the CRISPR/Cas9 machinery to the nucleus of the KGN cell line. Electroporator NEPA21 (Nepa Gene Co., Ltd., Chiba, Japan) was used to deliver pX458 plasmid containing specific guide RNAs and spCas9 endonuclease into KGN cells. After 48 hours, positive GFP cells were sorted and grew in normal culture conditions. Clone generation and genomic studies were performed as previously described.

### Western Blot analysis and proteomic study

Protein extraction and SDS-PAGE were performed according to standard protocols. Briefly, cells were lysed by RadioImmunoPrecipitation Assay (RIPA) buffer (150 mM NaCl, 0.1% SDS, 50mM Tris-HCL (pH 7.5), 0.5% sodium deoxycholate, 1% Nonyl Phenoxypolyethoxylethanol (NP-40) supplemented with protease and phosphatase inhibitors followed by protein dissociation using an ultrasonic processor UPH100H (Hielscher, Teltow, Germany) with 3 pulses of 5 seconds. Samples were centrifuged at 14,000 rpm at 4°C for 5 min and protein concentration was quantified following the BCA assay. Protein separation, transfer, antibody incubation, blocking and ECL detection were performed as indicated in Martin *et al*. [28]. Primary and secondary antibodies are detailed in Table 1.

Individual cell protein extracts (10 µg) were used for the TMT 10-plex quantitative proteomics analyses (see previous works for full protocol [29,30]. TMT experiments were analyzed on an Orbitrap Exploris 480 mass spectrometer (Thermo Fisher Scientific) equipped with the FAIMS Pro Duo interface (Thermo Fisher Scientific). Peptide separation was performed on the Vanquish Neo UHPLC System (Thermo Fisher Scientific) using established protocols [29]. MS data were analyzed with MaxQuant (version 2.1.3, Max Planck Institute of Biochemistry, Planegg, Germany) using standardized workflows. Mass spectra *.raw files were searched against the Uniprot UP000005640_9606.fasta Homo sapiens (human) 2022 database (20,577 protein entries) using reporter ion MS2 type for TMTs. Data normalization and data analysis was performed according to established protocols [29,31]. Proteins identified with one or more unique peptides, +/1.5 log2 foldchange values, and an adjusted p-value ≤ 0.05 were selected as statistically significant dysregulated proteins. Cutoffs were selected according to previous [29,30,32]. Analysis of the results was completed by GSEA and manually clusterization using Cytoscape (including EnrichmentMap and AutoAnnotate).

### RNA extraction and transcriptomic characterization

RNA processing involved TRIzol based isolation method followed by DNAse I treatment in all the cases. Sample quality was tested on an Agilent Bioanalyzer 2100 Eukaryote Total RNA nano assay. A whole-transcriptomic study was conducted by preparing total RNA libraries, as specified in the TruSeq Stranded Total RNA manufacturer instructions. Next steps involved pooling the samples in equimolar concentrations for sequencing on the NextSeq 500 instrument, preceded by various quality tests using the Agilent Bioanalyzer 2100 High Sensitivity DNA kit. Data obtained was analyzed by the Bioinformatic Unit of our center applying the same statistical parameters as in the proteomic study: +/1.5 log2 foldchange and padj<0.05. Enrichment in GO terms was conducted using preranked [33] GSEA software and network representation of enriched terms was obtained using enrichmentMap and AutoAnnotate from Cytoscape. Heatmaps and unsupervised clustering of top differentially expressed genes was generated using Morpheus and Volcano plots using VolcaNoseR.

For validation studies of the trancriptomic results, same RNA samples were subjected to RT-qPCR using SYBR green PCR master mix in the StepOne Real-Time PCR system and the appropriated primers (Table 1).

### 2.5. Cell growth, senescence, cell cycle migration and drug sensitivity

Cytell Cell Imaging System (GE Healthcare Life Sciences, Marlborough, MA, USA) was used as a digital microscope and cell counter in all the cell growth assays carried out along the study. For these experiments, 1500 cells from each clone were seeded in 96 well black glass bottom plates and maintained in culture under normal CO2 (5%) and temperature (37°C) conditions. At selected time points, cells were stained with 1µg/ml Hoechst and 10ug/ml of propidium iodide for 15-30 minutes at 37°. Cell viability was assessed using BioApp for image acquisition, and the generated data was analyzed using GraphPad software.

To analyze kinetic aspects of mitotic cell division, cells were previously infected with lentiviral particles expressing Histone 2B fused to green fluorescent protein and then seeded in a 6 well glass bottom plate. Images were captured every 30 minutes by fluorescence microscopy.

**D**rug sensitivity assays were performed using THUNDER Imager Live Cell & 3D Cell Culture & 3D Assay imaging system (Leica-Microsystems, Wetzlar, Germany). Cells treated with different concentrations of Ketoconazole, Dasatinib, Palbociclib and BRD6929 for 48 hours, were stained with Hoestch and propidium iodine (the same concentrations as above) and images of each experimental condition were taken to establish cell number and death rates. Drug reconstitution and dilution were carried out following datasheet instructions. Drug eligibility was based on previous published works [12,34,35] and Cmap (Connectivity Map, Broad Institute) results derived from our transcriptomic and proteomic studies. Data of differentially expressed genes (DEGs) with a +/1.5 log2 foldchange and a padj<0.05 and differentialy expressed proteins (DEPs) with a +/1.25 log2 foldchange and a padj<0.05 were compared and the resulting common genes were then inputted into the Cmap platform for further analysis. Finally, Palbociclib and BRD6929 were selected among the 3 major categories from a list of 100 drug candidates grouped by different description/activity features.

Senescence was determined using β-Galactosidase staining. A total of 70.000 cells per clone were seeded in 6 well transparent plates. Two days later, when cell confluency reached 60-70%, cells were fixed and stained following manufacturer’s recommendations (Senescence βgalactosidase Staining kit from Cell signaling). After cell staining, a total of 11 images comprising every part of the well were taken using Leica DM IL LED inverted Laboratory Microscope with the 10x objective and the Leica MC170 HD Microscope camera. Quantification was done by ImageJ software and results were represented as the percentage of cells exhibiting a positive β-galactosidase staining pattern.

A EdU incorporation assay was carried out to determine the percentage of cells that were at S-phase. Cells were seeded in glass cover slips placed in 6 well transparent plates. After reaching 50% confluency, EdU was added to the medium at a final concentration of 10 µM. One hour later, we proceed to perform Edu and nuclei staining according to manufacturer’s instructions. EdU labelling was detected by fluorescent microscopy (Leica-Microsystems, Wetzlar, Germany) employing 10x and 20x objectives (N PLAN 10x/0.25 DRY and HC PL FLUOTAR 20x/0.55 DRY). Percentage of positive EdU cells were determined by ImageJ software.

The invasive capacity was analyzed by a Transwell invasion assay as described in Martin et al.,2023 [28], with minimal modifications. In brief, once 2000 cells were seeded in low serum conditions in the upper part of the chamber, we waited for 48 hours before removing Matrigel from the lower membrane where cells are attracted by the 10% serum containing media placed in the bottom well. Next, cells were fixed and stained with 1% glutaraldehyde and 0.5% crystal violet, respectively. Images were taken with a Leica MC170 HD Microscope camera coupled with a Leica DM IL LED inverted Laboratory Microscope at 10x magnification. Number of invasive cells was determined by ImageJ software.

### 2.6. Zebrafish care and handling. Xenograft assays and image analysis

Zebrafish adult fish (*Danio rerio*, wild type) were crossed to obtain zebrafish embryos. Adult zebrafish were kept in aquaria with a ratio of 1 fish per liter of water, a day/night cycle of 14:10h and water temperature of 28’5°C was constantly maintained, according to the published protocols [36]. Procedures used in the experiments, fish care and treatment were performed in agreement with the Animal Care and Use Committee of the University of Santiago de Compostela and the standard protocols of Spain (Directive 2012-63-DaUE). At the end of the experiments, the embryos used were euthanized by tricaine overdose.

Zebrafish embryos were collected at 0hpf (hours post fertilization) and incubated at 28.5°C in petri dishes until 48hpf. Then, KGN cells were trypsinized and concentrated in an Eppendorf at a rate of ≈1 million cells for each condition and resuspended in 10µL of PBS (phosphate buffered saline) with 2% of PVP (Polyvinylpyrrolidone) to avoid cellular aggregation. After cell preparation, 48hpf embryos were anesthetized with 0.003% of tricaine (Sigma) and injected with the corresponding clones. Cell injection was performed using borosilicate needles (1 mm O.D. x 0.75 mm I.D.; World Precision Instruments). Between 100 and 200 cells were injected into the circulation of each fish (Duct of Cuvier) using a microinjector (IM-31 Electric Microinjector, Narishige) with an output pressure of 20kPA and 15ms of injection time per injection. Afterwards, embryos were incubated until 6 days post injection (dpi) at 34°C in 30mL Petri dishes with SDTW (Salt Dechlorinate Tap Water). Imaging of the injected embryos were performed using a fluorescence stereomicroscope (AZ-100, Nikon) at 1, 4 and 6dpi to measure the spreading and proliferation of the injected cells in the caudal hematopoietic tissue (CHT) of the zebrafish embryos. To perform the analysis of the acquired images at the different time points Quantifish software [37] was used. This software processes all the images and performs the measure of the fluorescence intensity and area of positive pixels, corresponding to the cells injected, above a certain threshold. With these parameters, integrated density is obtained allowing the researcher to compare different times between images and obtain a proliferation ratio of the cells in the region of the caudal hematopoietic tissue (CHT) of the embryos, where the cells normally metastasize. As a measure of the metastatic capacity of the injected cells, the number of tumors present in the tails of the embryos were scored. Tumor counts were determined based on the size of the tumor and the separation between the cell clusters inside the tail of the fish.

### 2.7. Statistical analyses

Graphpad software was used to perform the statistical analysis of all the data. ANOVA test and Student’s t-test were used to compare edited clones to their reference controls (parental cells) with a confidence interval of 95%.

## Results

### Precise disruption of FOXL2 c.402C>G, p.C134W mutation via CRISPR/SpCas9 genome editing system

To study the role of FOXL2-C134W mutation in tumor maintenance, an allele-specific gene editing approach based on CRISPR technology was developed, to selectively target the mutant allele without affecting its corresponding wild-type counterpart. This last aspect is especially relevant since the indiscriminate destruction of both alleles might cause side effects. We hypothesized that the FOXL2 c.402C>G, p.C134W allele could be specifically inactivated by exploiting the fact that this pathogenic missense mutation gives rise to a nucleotide variation in the seed sequence (one to five nucleotide upstream of the PAM) that potentially affect SpCas9 recognition. Following this strategy, we designed two allelespecific sgRNA, sg1.3 and sg1.4, that include the c.402C>G mutation at position +7 and +2, respectively, upstream of the PAM (Figure 1a). The specificity of these guides was assayed in KGN cells, the only commercially available cell line derived from an ovarian granulosa human tumor that harbors the FOXL2 c.402C>G mutation in heterozygosity. As described in Figure 1b, CRISPR machinery was sequentially introduced in the target cells by lentiviral infection resulting in three pool cell populations expressing Cas9 alone or in combination with sg1.3 or sg1.4 small guide RNAs.

**Figure 1.**
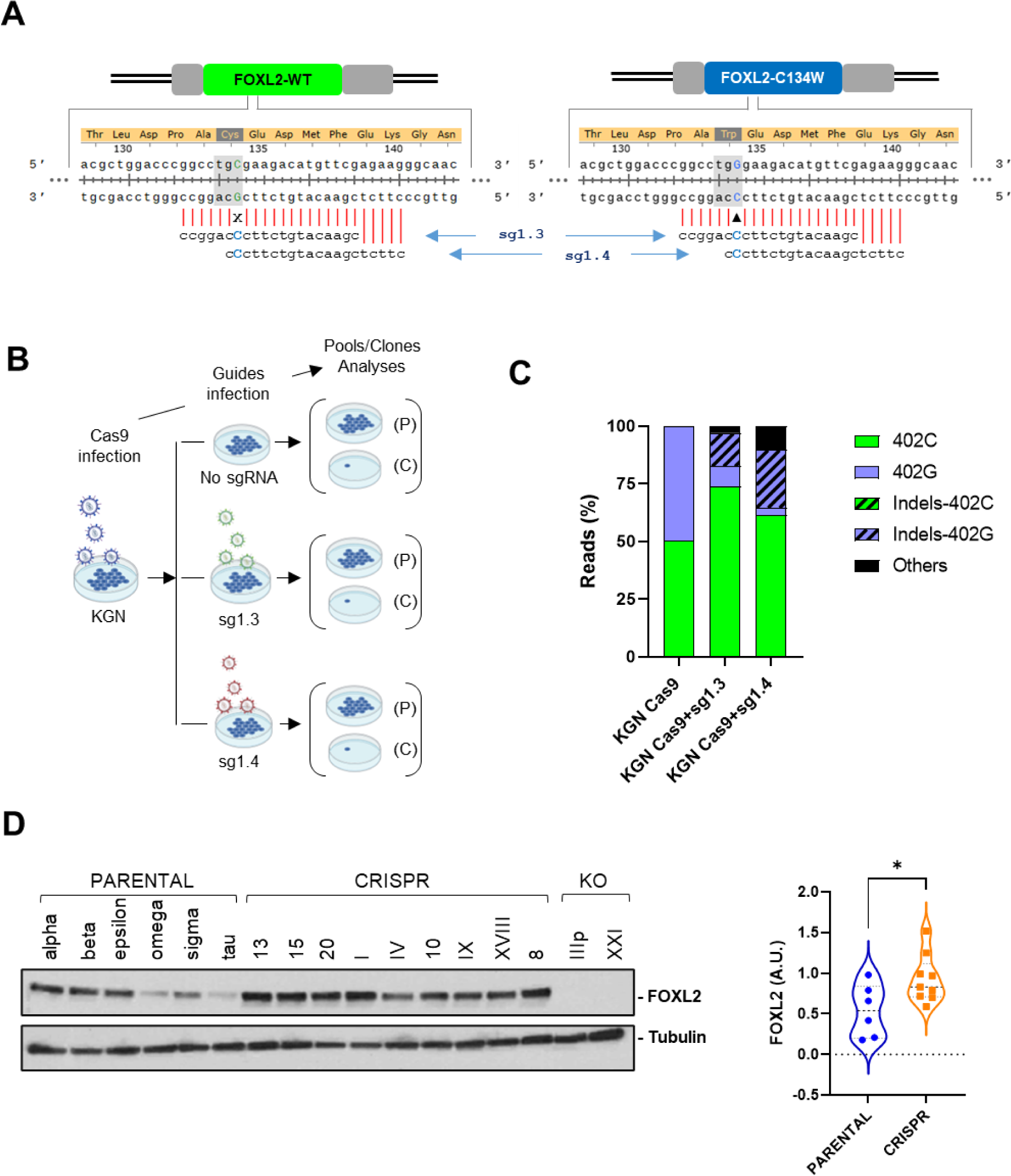
**Edition of FOXL2 c.402C>G mutation in KGN cells**. **(A)** Scheme for FOXL2 c.402C>G sgRNA location. FOXL2 wild type and mutant alleles are represented along with the corresponding aminoacid sequences for both alleles. Mutant and wild type bases are highlighted in blue and green, respectively. Guides designed specifically to target mutant allele are represented below each allele. **(B)** Timeline for the generation of pools (P) and clones (C). Each round of infection is followed by a selection performed with antibiotics (Blasticidin and Puromycin, respectively). **(C)** Guide specificity on the FOXL2 c.402C>G mutant allele. Graph represents percentage of reads matching each edited and not edited alleles for the pool of cells generated upon infection with Cas9, Cas9+sg1.3 and Cas9+sg1.4. **(D)** FOXL2 protein expression in parental, CRISPR and KO clones isolated in this study. Violin graph shows the quantification of FOXL2 expression in parental and CRISPR clones.

Subsequent genomic analysis by deep amplicon sequencing allowed us to potentially distinguish between 5 different allelic variants (Figure 1c). Two, non-edited, consist of the wild type and mutant alleles (referred as 402C and 402G, respectively). The other two are the modified wild type and mutant alleles containing several types of insertions and deletions (designated as Indels-402C and Indels-402G, respectively). In a fifth category (named as others), we could not identify which of the two alleles had been edited. As expected, we only detected, at approximately 50% frequency, wild type (402C) and mutant (402G) alleles in the pool expressing the SpCas9 alone (KGN-Cas9) (Figure 1c; Supplementary Table 1). Importantly, the co-expression of Cas9 with any of the two sgRNAs caused a dramatic reduction in the representation of the mutant, 402G allele. In the case of the sg1.3 guide, we could only detect 8.85% of the intact 402G allele, while the guide sg1.4 was associated with the presence of just 3.44% 402G reads. As expected, this reduction of 402G alleles was accompanied by a corresponding increase of Indel-402G alleles, which were detected in 14.31% and 25.04% of the reads from the sg1.3 and sg1.4 infected cells, respectively. Additional modifications (others) were also identified in 3.02% and 10.27% of the reads from the corresponding sg1.3 and sg1.4 infected cells. It is interesting to note, the increase of intact, wild type reads detected in sg1.3 and sg1.4 infected cells when compared with the control, KGN-Cas9 ones (73.85% and 61.24%, respectively, vs 50.21%). Finally, a specific pattern of indels is detected in a sgRNA dependent manner. While the most common indel in the pool containing sg1.3 is the deletion of a cytosine in position 399, the insertion of an adenine between positions 403 and 404 is the most frequent indel promoted in the case of sg1.4 (Supplementary Table2). Similar results were obtained when the CRISPR complexes were introduced in KGN cells by nucleofection (Supplementary Figure 1).

In conclusion, although a complete absence of genomic edition in the wild type allele could not be unambiguously demonstrated, the inability to identify the presence of genetic modifications, unlike in the case of the mutant, 402G allele, underlines the selectivity of both sg1.3 and sg1.4 guides towards the Cas9-dependent disruption of the mutated FOXL2 allele.

To further analyze the consequences of CRISPR-mediated elimination of the FOXL2 c.402C>G mutation, we isolated and characterized individual clones from the pools transfected with the different CRISPR complexes. A total of 37 clones obtained by a single cell cloning approach were characterized by sequencing. As expected, all the 10 clones derived from the cell pool that was transfected with SpCas9 without sgRNAs, are identical to the original KGN cells (data not shown), which means that they preserve an intact FOXL2 mutation in heterozygosity. These PARENTAL clones were identified with Greek letters and will be used as controls for later analyses. Remarkably, the majority of the remaining clones (25 out of the 27 isolated from pools co-expressing SpCas9 and either of the two allele-specific sgRNAs, hereafter referred as CRISPR clones) showed some type of CRISPR activity on the mutant allele (Supplementary Table 3). Most of those cases (21 out of 25) displayed Indel events that exclusively occurred on the mutant allele with frame shift modifications leading to premature stop codons (Supplementary Table 3). The observed Indel pattern was like the one already detected in the pools (single cytosine deletion at position 399 in the case of the sg1.3 clones and a prevalent adenine insertion between nucleotides 403 and 404 for the sg1.4 ones (Supplementary Table 3). The guide origin of the clones is reflected in their nomenclature with Arabic and Roman numbers to designate clones coming from sg1.3 and sg1.4, respectively. Although small insertions or deletions are the major causes of allele-specific inactivation (77,8%), we failed to detect the presence of the mutant allele in a small percentage of clones (4 out of 27, 14,8%), indicating that a large deletion occurred at the mutant allele or, alternatively but less probable, DNA correction by homology-directed repair took place in the cell using the wild type allele on the homologous chromosome as a template. These 4 clones were also included in the CRISPR group. Finally, two knockout (KO) clones (IIIp and XXI) harbor indels in both alleles that provoke changes in the open reading frame that result in truncated, and presumably nonfunctional, wild type and mutant proteins (Supplementary Table 3). Therefore, this finding suggests that, although to a much lesser extent, biallelic modification by CRISPR can also occur in our experimental system.

To complete the molecular characterization of these clones, we analyzed by western blot the impact of CRISPR mediated genome edition of *FOXL2* on its protein expression. Interestingly, while FOXL2 levels were completely abrogated in the two KO clones, CRISPR clones showed a significant FOXL2 upregulation compared to parental KGN controls (Figure 1d). Moreover, specific analysis of wild type FOXL2 mRNA by RT-qPCR, revealed a clear increase when mutant allele is selectively inactivated (Supplementary Figure 2), suggesting some kind of transcriptional repression mechanism operating in the presence of the FOXL2-C134W mutation. To further explore this possibility, we analyzed also the expression of CYP19A1, a well-known target of FOXL2 [38]. As expected, elevated levels of CYP19A1 were detected in CRISPR clones, which reverted the almost complete inhibition of this gene characteristic of the PARENTAL, KGN cells (Supplementary Figure 2).

**Figure 2.**
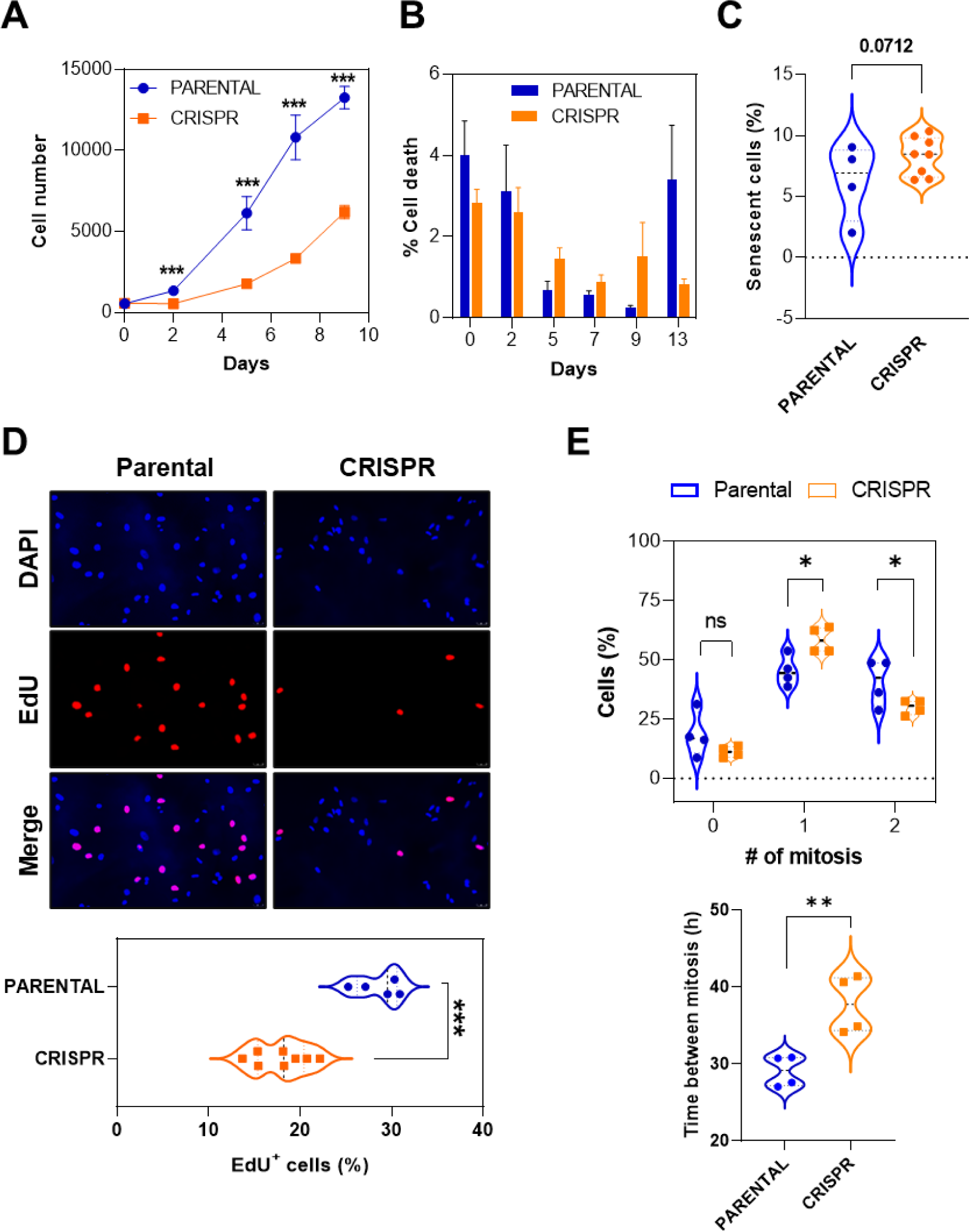
Specific FOXL2 c.402C>G mutation elimination causes a decrease in cell growth. (A) Growth curves of PARENTAL (n=5) and CRISPR (n=9) clones. Each point corresponds to the average of 3 different experiments. Bars represent standard error of the mean (SEM). **(B)** Percentage of cell death related to each day of the growth curve showed in panel (a). No significant differences were found after the analysis between the different groups. **(C)** Graph represents mean of percentage of positive cells for βGalactosidase staining in PARENTAL (n=4) and CRISPR (n=8) clones. **(D)** Cells proliferation in PARENTAL (n= 5) and CRISPR (n=8) clones estimated by incorporation of EdU. The upper panel corresponds to representative clone images from both groups, showing EdU incorporation (red, middle images) and all nuclei (blue, upper images). Bottom graph shows percentage of cells positive for EdU staining in the nucleus. **(E)** CRISPR clones show modifications in cell cycle timing. The upper graph shows the percentage of cells that undergo 0, 1 or 2 rounds of mitosis for 48 hours. Bottom violin graph shows the quantification of the time lapse between two subsequent mitoses in CRISPR and PARENTAL clones. In all the experiments of this figure, data was compared with one-tailed, unpaired Student’s t-test (*p<0.05; **p<0.01; ***p <0.001).

In conclusion, we have generated a CRIPSR-mediated system that allows us to specifically eliminate the FOXL2 c.402C>G allele in KGN granulosa tumor cells. Moreover, the abrogation of the FOXL2-C134W mutant restores wild type function of FOXL2 as a transcription factor.

### *In vitro* targeted disruption of FOXL2 c.402C>G allele diminishes cell proliferation of AGCT derived cells

To investigate whether specific cleavage of FOXL2 c.402C>G mutant allele mediated by CRISPR could induce proliferative defects, we initially conducted viability assays of parental and CRISPR clones. Results show that proliferation capacity of edited cells was diminished compared to non-edited controls (Figure 2a). In the same experiments, cell death levels were not able to explain this antiproliferative phenotype since no significant differences were detected between CRISPR and parental clones (Figure 2b). Interestingly, in a close to significance way, the percentage of senescent cells was increased among CRISPR clones when compared with the parental ones (Figure 2c). We then further assessed the cell cycle of both clonal populations finding a reduced capacity of sg1.3and sg1.4targeted cells to incorporate EdU which indicates a potential delay in S-phase entry and/or retarded progression (Figure 2d).

Next, we performed time-lapse microscopy analysis to monitor in more detail cell cycle kinetics by imaging for 48 hours H2B-EGFP expressing cells of both genotypes. Data indicate that, although no differences were detected for the percentage of cells that do not undergo mitosis, most CRISPR cells tend to divide only once, whereas a great fraction of PARENTAL cells exhibit two rounds of cell division in the same period (Figure 2e). Confirming these results, CRISPR clones needed significantly more time to go from one mitosis to the next one than the PARENTAL ones (38±4h vs 29±2h). To try to decipher the molecular mechanism responsible for this reduced proliferative ability associated with FOXL2C134W inactivation, different signaling pathways involved in the regulation of cell survival and proliferation were analyzed. While no differences were observed in terms of ERK1/2 proteins, phosphorylation of AKT, GSK3β, and Smad2/3 phosphorylation was significantly reduced in CRISPR clones (Figure 3a). Interestingly, this phenomenon was accompanied by a significant increase of p27 (Figure 3b), a FOXL2 regulated player [39,40] involved in cell cycle inhibition whose expression is known to be also negatively controlled by PI3K pathway [41].

**Figure 3.**
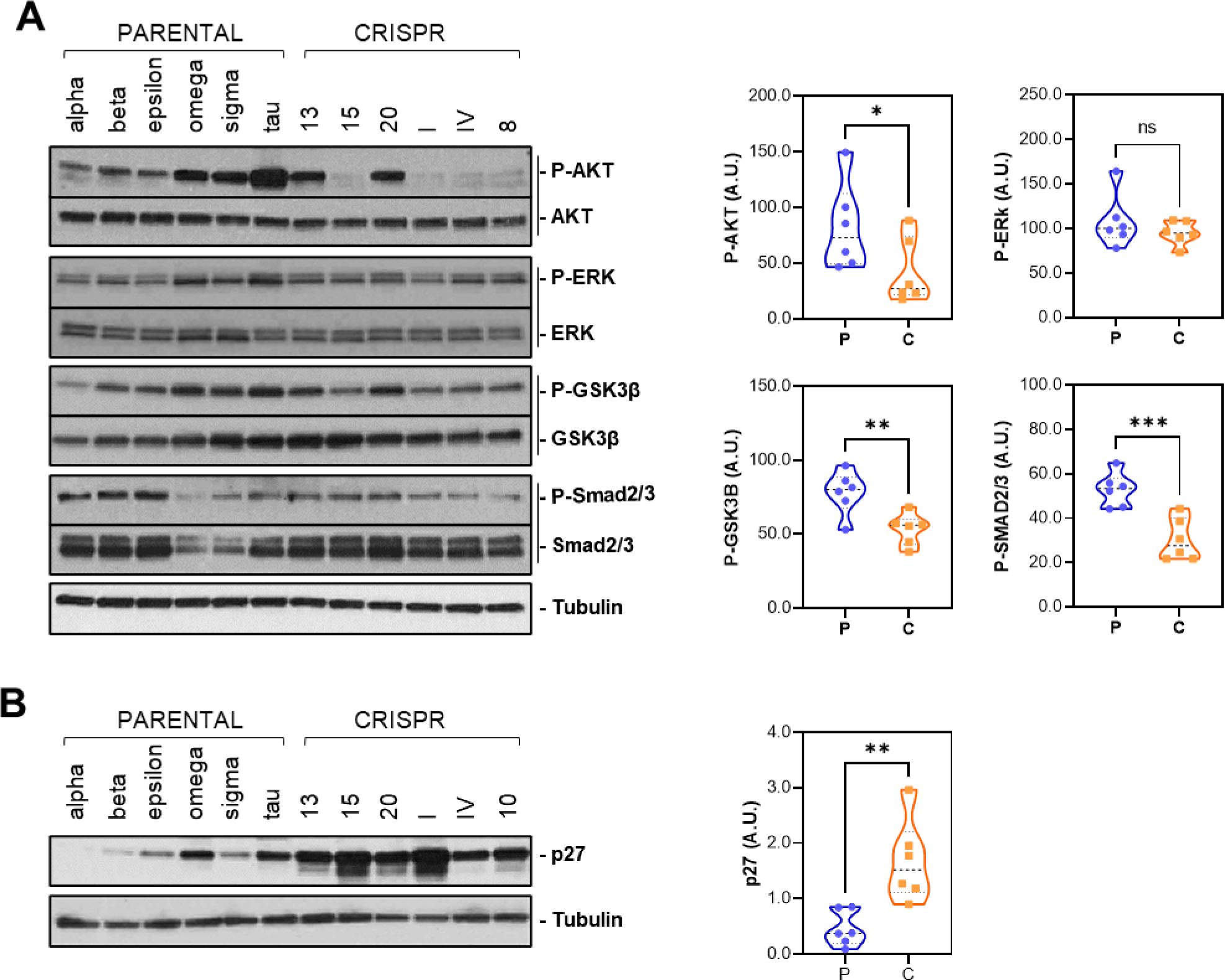
Cell survival and growth regulation pathways are affected by the specific elimination of FOXL2 c.402C>G mutation in KGN cells. Protein expression of molecules involved in proliferative pathways **(A)** and p27 **(B)**. Six different PARENTAL and CRISPR clones are analyzed. Right graphs show quantification of the expression results showed in (A) for phospho-proteins and in (B). Each phospho-protein was normalized with its total expression and the expression of tubulin. Proteins in B were normalized with tubulin protein levels.

Altogether, these data let us conclude that specific abrogation of FOXL2 C134W mutation negatively impact on cell proliferation by causing a deceleration in the cell cycle and a moderate increase in senescence. These changes in cell fate are associated to inhibition of PI3K/AKT, and TGFβ-Smad2/3 pathways as well as the concomitant upregulation of the cell cycle inhibitor p27.

### Specific elimination of mutated FOXL2 allele attenuates cell migration both *in vitro* and *in vivo* and predispose to a better response to drug therapies

Most of the AGCT clinical cases do not exceed stage I at the time of diagnosis and have good prognosis. However, a small percentage of these patients can relapse, showing metastatic lesions in different locations that correlate with poor survival rates [42,43]. Thus, we aimed to analyze the invasive potential of both clonal groups by standard transwell analysis. We seeded the cells on top of the filter membrane of the transwell insert and exposed the cells to a chemoattractant that was added on the bottom part of the well. Forty-eight hours later, as shown in Figure 4a, CRISPR clones showed less invasive capacity than the PARENTAL ones, indicating that this malignant feature was also diminished upon the elimination of the FOXL2 c.402C>G mutation.

**Figure 4.**
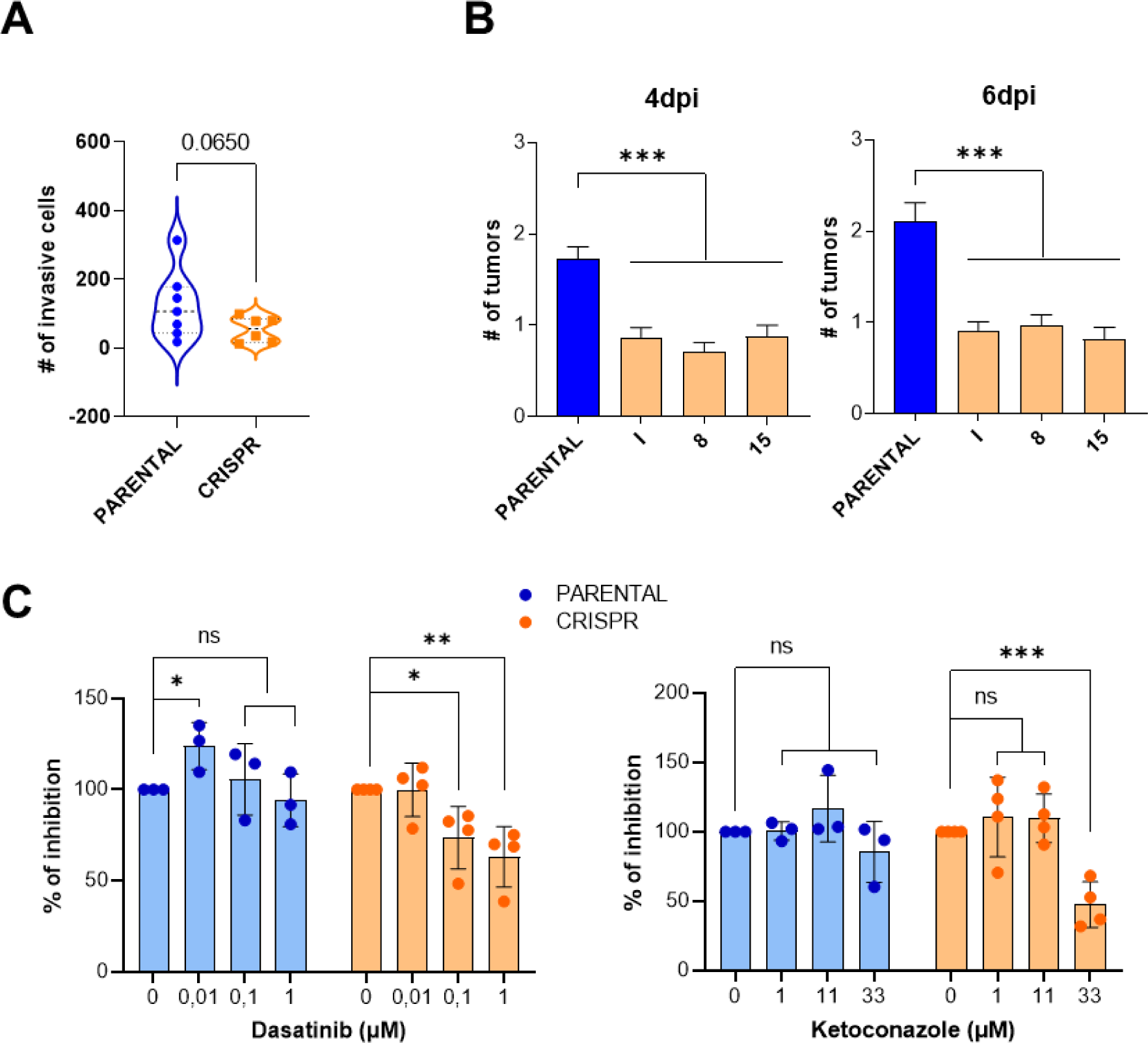
Migration capacity and sensitivity to Dasatinib and Ketokonazole is altered in CRISPR clones. (A) *In vitro* migration assay. Violin plots show all the clones individually analyzed for the experiment, 48 hours after seeding cells in transwells. A close-to-significance increase in the average number of invasive cells is found among CRISPR clones when compared to PARENTAL ones. **(B)** *In vivo* migration assay in zebrafish. Graphs represent the number of tumors counted in animal tail. PARENTAL clones are unified in a single (blue) column while CRISPR clones (orange) are shown individually. Statistical differences are observed at 4 and 6 days post cells injection (4dpi and 6dpi, respectively), revealing a reduction in tumor formation in CRISPR clones. **(C)** CRISPR-mediated elimination of FOXL2 4c.02C>G mutation and drug therapy have a synergistic effect in cell inhibition. Percentage of cells with different doses of Dasatinib (left) and Ketoconazole (right) 48 hours post drug addition. Each cell number value was normalized taking as 100% the number of cells in control wells (0 µM concentration for each drug tested). No significant growth inhibition was shown in PARENTAL clones, while cell number was significantly reduced in CRISPR clones treated with Datasitinb or Ketoconazole. Statistical analyses were carried out in each group comparing different drug doses with the untreated condition. (* p<0.05; ** p<0.01, *** p<<0.001).

Considering that *in vivo* studies in zebrafish have been reported to faithfully determine the metastatic potential of a broad range of cancer cells [44], two clones per each FOXL2 genotype were injected into the circulation of 48 hour-postfertilization embryos and incubated at 34°C for 6 days. The number of micrometastasis was quantified in the embryo tails by taking images at 1, 4 and 6 days post-injection as a measurement of the spread and metastatic capacity of FOXL2 targeted and non-targeted KGN cells. As illustrated in Figure 4b, a significant reduction of around 50% in tumor burden was scored when CRISPR clones were injected compared to PARENTAL ones, suggesting an important role of the FOXL2 C134W mutation in the dissemination ability of AGCT derived cells.

Although treatment of AGCTs is mainly based in surgery and platinum-based combinations, a progressive characterization of these tumors along the years have allowed to introduce and propose new therapeutic approaches. The senolytic Dasatinib and the CYP17 inhibitor Ketoconazole are among these alternatives. Dasatinib has showed a synergistic effect when used with chemotherapy in AGCTs [34]. On the other hand, inhibition of CYP17 has been proposed as a promising therapeutic option for AGCT [12] because FOXL2 c.402C>G mutation leads to overexpression of this key regulator of steroidogenesis [45]. In fact, Ketoconazole, a CYP17 inhibitor, has shown activity in advanced AGCT in clinical and *in vitro* studies [35]. Therefore, we decided to study the activity of these two compounds in a contest of CRISPR-mediated elimination of the FOXL2 c.402C>G mutation. As shown in Figure 4c, cell growth is significantly inhibited in CRISPR clones treated with either Dasatinib or Ketoconozale, whereas no inhibitory effect is detected in PARENTAL clones treated with the same drugs.

### Elimination of FOXL2 c.402C>G mutation modifies the transcriptomic expression signature of granulosa tumor cells

It has been previously described that mutation 402C>G in FOXL2 transcription factor, due to its location in the DNAbinding domain, affects the expression of several target genes [26], changing the transcriptomic profile of granulosa cells. It has been demonstrated, both *in vitro* and *in vivo*, that these alterations affect pathways related with TGFβ signaling, ECM organization, ovarian infertility genes, regulation of gene expression by several interleukins, FSH apoptosis, cholesterol and prostaglandin biosynthesis and regulation [46,47]. To gain insights into the transcriptional effects of eliminating the 402C>G mutation of FOXL2 in KGN cells, we compared the expression signatures of CRISPR clones (n=3) versus those of the PARENTAL ones (n=4). Principal component analysis (PCA) (Figure 5a) of the different clones in the experiment revealed a clear separation between the PARENTAL and CRISPR clones, defining them as distinct entities. Differentially expressed genes (DEG; fold change>1.5 or <1.5 and adjusted p-value of <0.05) in PARENTAL clones, when compared with CRISPR counterparts, include a total of 710 upregulated and 943 downregulated transcripts (Figure 5b; Supplementary Table 4). These results were validated by RT-qPCR of a selected group of DEGs (Supplementary Figure 2).

**Figure 5.**
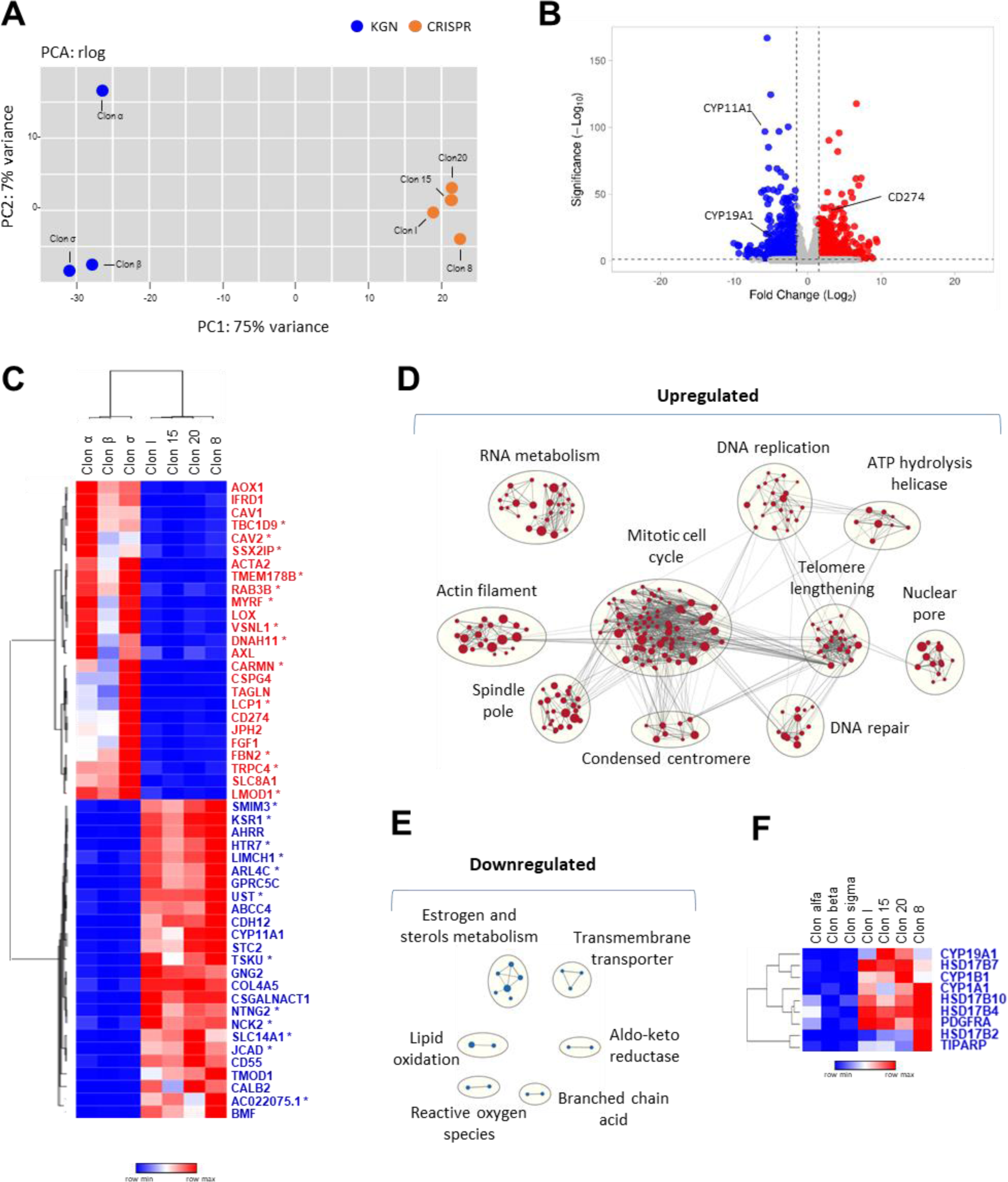
CRISPR-mediated elimination of FOXL2 c.402C>G mutation induces significant changes in the transcriptomic profile of granulosa tumor cells. (A) Principal component analysis of RNAseq data from KGN (blue) and CRISPR (red) clones. **(B)** Volcano plot for differential gene expression in FOXL2 C134W (KGN) vs edited (CRISPR) clones. Downregulated genes with adjusted Pvalue<0.05 and log2 of fold change lower than -1.5 are depicted in blue. Upregulated genes with adjusted P-value<0.05 and log2 of fold change higher than 1.5 are depicted in red. **(C)** Heatmap showing an unsupervised clustering of top 25 upregulated and downregulated genes according to adjusted p-value. Genes that have not been related to granulosa cells are marked with an asterisk. **(D)** Clusters of GO gene sets differentially enriched in genes upregulated in C134W vs wild type (CRISPR) clones at q-value<0.05. Node size is proportional to the number of genes identified in each gene set. The light grey edges indicate gene overlap between gene sets. **(E)** Clusters of GO terms at p-value<0.05 enriched in downregulated genes depicted as in D. **(F)** Heatmat of genes belonging to the GOBP estrogen metabolic process significantly downregulated in KGN clones at adjusted p-value>0.05.

Most of the top-50 most up/downregulated DEGs (Figure 5c) have been previously related with cancer at some extent (Supplementary table 5). Some of them (CAV1, CAV2, KSR1) are involved in MAPK signaling, whereas others have been associated with TGFβ signaling (LOX, TSKU), extracellular functions/junctions (JCAD, COL4A5, CDH12) and apoptosis (BMF), among other cellular features. It is noteworthy that PDL1 (CD274) ranks among the top 25 most upregulated genes in clones harboring the C134W mutation. This upregulation could potentially promote immune response evasion, thereby facilitating tumor growth. However, overexpression of PDL1 in FOXL2 C134W tumors might render these tumors more sensitive to anti-PDL1 therapy. Furthermore, 50% of the top-50 most up/downregulated DEGs have already been associated with granulosa cells and/or FOXL2 (Figure 5c; Supplementary Table 5). Hence, CSPG4, JPH2 and TAGLN have also been found upregulated in TGFβ -induced GCTs, when compared with wild type granulosa cells [48]. On the other hand, among the most significantly downregulated genes, CDH12 and CD55 are granulosa cell markers [49,50], whereas CYP11A1, a FOXL2 target gene [51], plays a crucial role in steroid hormone synthesis in granulosa cells [52].

To further explore the potential roles of the DEGs, enrichment of GO terms was analyzed using GSEA. As shown in Figure 5d, up-regulated genes in PARENTAL vs CRISPR clones were enriched in GO gene sets related to cell cycle, including DNA replication and repair, spindle pole, mitosis and centromere condensation. These results correlate with the differences observed between the two clone groups in terms of cell proliferation and mitosis progression and duration (Figure 2). Upregulated genes were also significantly enriched in pathways related to cell cycle, DNA repair and others linked to protein synthesis and post-translational modifications like sumoylation (data not shown). Further, several gene sets related to estrogen metabolism, a critical function of granulosa cells [53,54] where FOXL2 have been previously implicated [25,55], were downregulated in C134W cells compared to wild type CRISPR clones(Figure 5e, 5f). All this transcriptomic information indicates that the CRISPR-mediated elimination of the C134W mutation leads to the acquisition of a different cellular status. To explore more this new condition, we decided to compare the transcriptomic profile of our PARENTAL and CRISPR clones with that generated previously by Weis-Banke and coworkers for human granulosa cells expressing different amounts and types of exogenous FOXL2 variants [26]. A hierarchical cluster analysis shows that CRISPR clones grouped close to human normal granulosa cells transfected with either empty, FOXL2wt or FOXL2-C134W expression vectors, whereas PARENTAL clones cluster closer to the KGN cells used by Weis-Banke *et al*. [56] (Supplementary Figure 3). These results indicate a potential reversion of KGN cells to their wild-type origin upon elimination of the C134W mutation.

### Protein expression signature of KGN cells is significantly altered upon the elimination of FOXL2 c.402C>G mutation and can be used to discover molecules with growth inhibitory activity against AGCT cells

To gain further insights in the expression profile associated with the lack of FOXL2-C134W mutation, we carried out a proteomic study comparing PARENTAL and CRISPR clones. Mirroring the transcriptomic results, two very differentiated groups appear upon PCA examination (Figure 6a). To identify differentially expressed proteins (DEPs), we filter our results by an adjusted p-value of <0.05 and a fold change beyond +/-1.5. Using these parameters, we detected a total of 29 downregulated and 60 upregulated proteins in PARENTAL clones when compared with the CRISPR group (Figure 6b; Supplementary Table 6). A Significant fraction of these proteins was also found de-regulated in the transcriptomic study (Figure 6c). Hence, 8 out of the 29 (27.6%) down-regulated proteins and 36 out of the 60 (65.6%) upregulated ones were also differentially expressed at the RNA level (Supplementary Table 7). Interestingly, a very good correlation was observed between RNA and protein expression of these 44 genes (Figure 6c). Likewise for the transcriptomic results, the top-50 most up/down DEPs clusterized the two clone types, CRISPR and PARENTAL, as two differentiated entities (Figure 6d). Most of these top-50 DEPs have already been associated with cancer processes and more than half of them (30 out of 50) have been related to granulosa cell function and/or FOXL2 (Supplementary Table 8). Interestingly, 8 out of the top-50 DEPs were also found in the RNA-DEGs top-50 list (Figure 6b and d). The upregulated ones include TAGLN, an actin-associated protein related with cytoskeleton organization; LMOD1, which is involved in follicle maturation; CSPG4, a regulator of cell-substratum interactions; and CAV1 and CAV2, which are main components of the cavolae plasma membranes.

**Figure 6.**
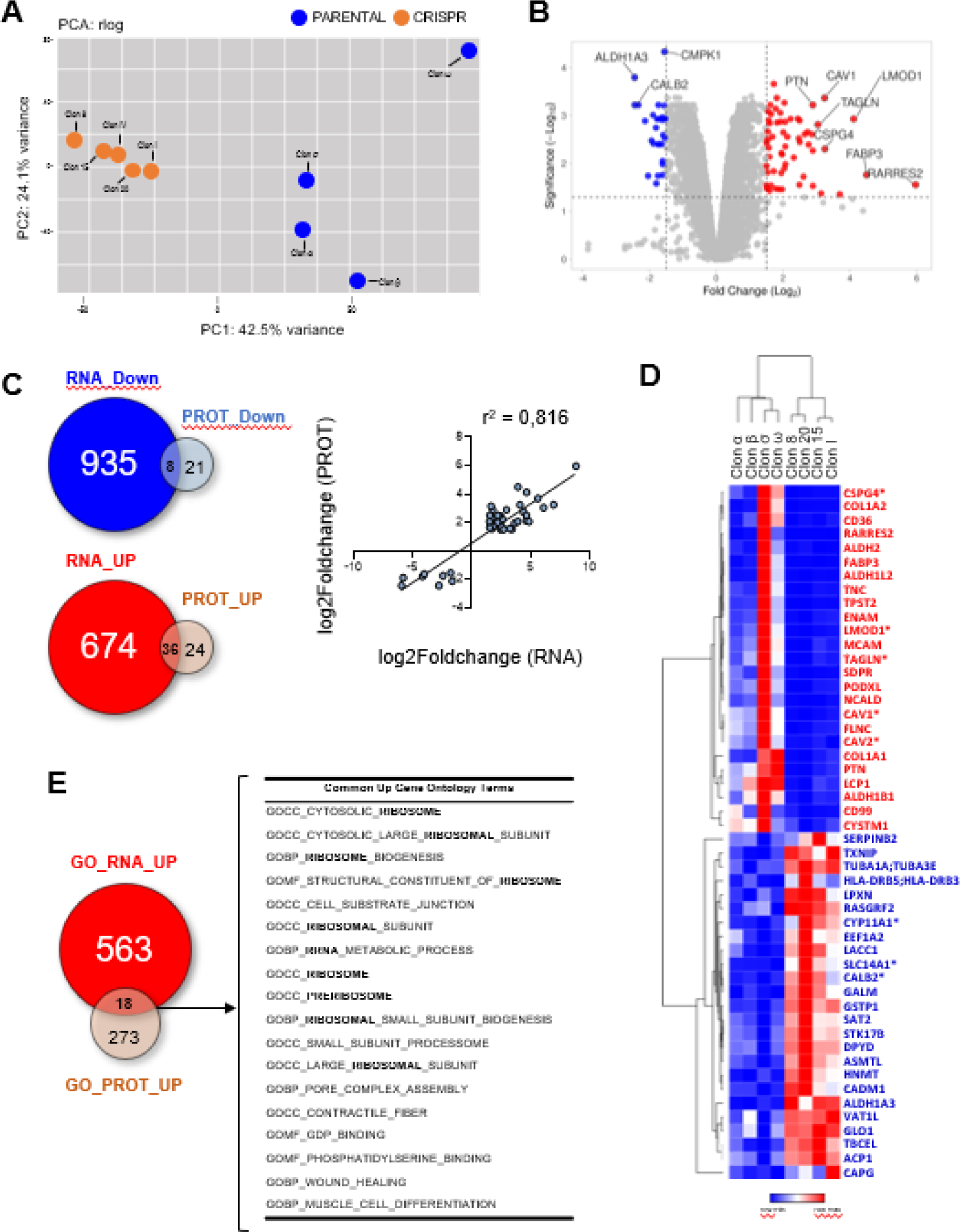
Proteomic changes induced by the CRISPR-mediated elimination of FOXL2 c.402C>G mutation. (A) Principal component analysis of protein data from KGN (blue) and CRISPR (red) clones. **(B)** Volcano plot for differential gene expression in FOXL2 C134W (KGN) vs edited (CRISPR) clones. Downregulated proteins with adjusted P-value<0.05 and log2 of fold change lower than -1.5 are depicted in blue. Upregulated genes with adjusted P-value<0.05 and log2 of fold change higher than 1.5 are depicted in red. **(C)** Venn-diagrams show the overlap of significantly de-regulated genes at the RNA and protein levels. Right plot shows the correlation between the RNA and protein fold change values of the common cases. **(D)** Heatmap of unsupervised clustering of the top 25 upregulated and downregulated proteins according to adjusted p-value. Genes that are also in the top-50 de-regulated list of the transcriptomic analysis are marked with an asterisk and indicated in panel b. **(E)** Venn diagram shows the GO terms at p-value<0.05 enriched in upregulated genes. Common GO terms are also listed in the adjacent right table.

Regarding the three commonly down-regulated, top-50 DEGs/DEPs, CALB2 plays a key role in calcium mediated signaling, SLC14A1 is a mediator of urea transport and CYP11A1 is a cytochrome P450 monooxygenase that catalyzes the side-chain hydroxylation and cleavage of cholesterol to pregnenolone, the precursor of most steroid hormones.

A GO analysis was also carried out with the DEPs found when PARENTAL clones were compared with CRISPR ones. For both up-regulated and down-regulated proteins, the GO terms detected significantly differed from the ones observed in the transcriptomic analyses (Supplementary Figure 4). GO-upregulated terms included “adhesion and extracellular structure”, “mitochondrial translation complex” and “cholesterol metabolic regulation” among others, whereas “Pyrimidine ribonucleotide biosynthesis”, “Toll receptor signaling” and “Multivesicular body” are among the GOdownreguated ones. It is interesting to note that GO terms common for both the transcriptomic and proteomic GO analyses were only detected for up-regulated genes and are mostly associated with ribosomes (Figure 6e).

In a final step of our work, we tried to identify compounds that can induce the same expression signature that characterizes CRISPR clones when compared to the PARENTAL ones. To identify such compounds, we used the commonly downregulated genes from our transcriptomic and proteomic analyses (Figure 7a) to interrogate the Connetivity Map (cMap) data base. This platform allows to search for similarities to a given expression signature among expression signatures induced in 9 cancer cell lines by more than 5,000 small molecules [57]. This *in silico* approach allowed us to rank these drugs according to cMap connectivity scores (CS) that predicts the similarity of the profile induced by each compound with the interrogated expression profile (Supplementary Table 9). Confirming the therapeutic value of this bioinformatic tool, Ketoconozale, an orphan drug used for the treatment of AGCT [12] ranks #59 among the list of compounds. Since the top-100 drugs were enriched in Histone deacetylase (HDAC), Topoisomerase and CDK inhibitors (Supplementary Table 10), we selected one HDAC inhibitor (Merck60 or BRD6929, ranked #63) and a CDK inhibitor (Palbociclib, #17 of the ranking list) to study their activity on PARENTAL cells. As shown in Figure 7b, Palbociclib was able to induce cell death at the highest dose it was used (10µM), whereas BRD6929 induced the same effect at all the doses we tested (0.01, 0.1, 1 and 10 µM). Furthermore, a significant reduction of the number of cells in culture when compared with untreated cells was observed at all the concentrations for both compounds (Figure 7c).

**Figure 7.**
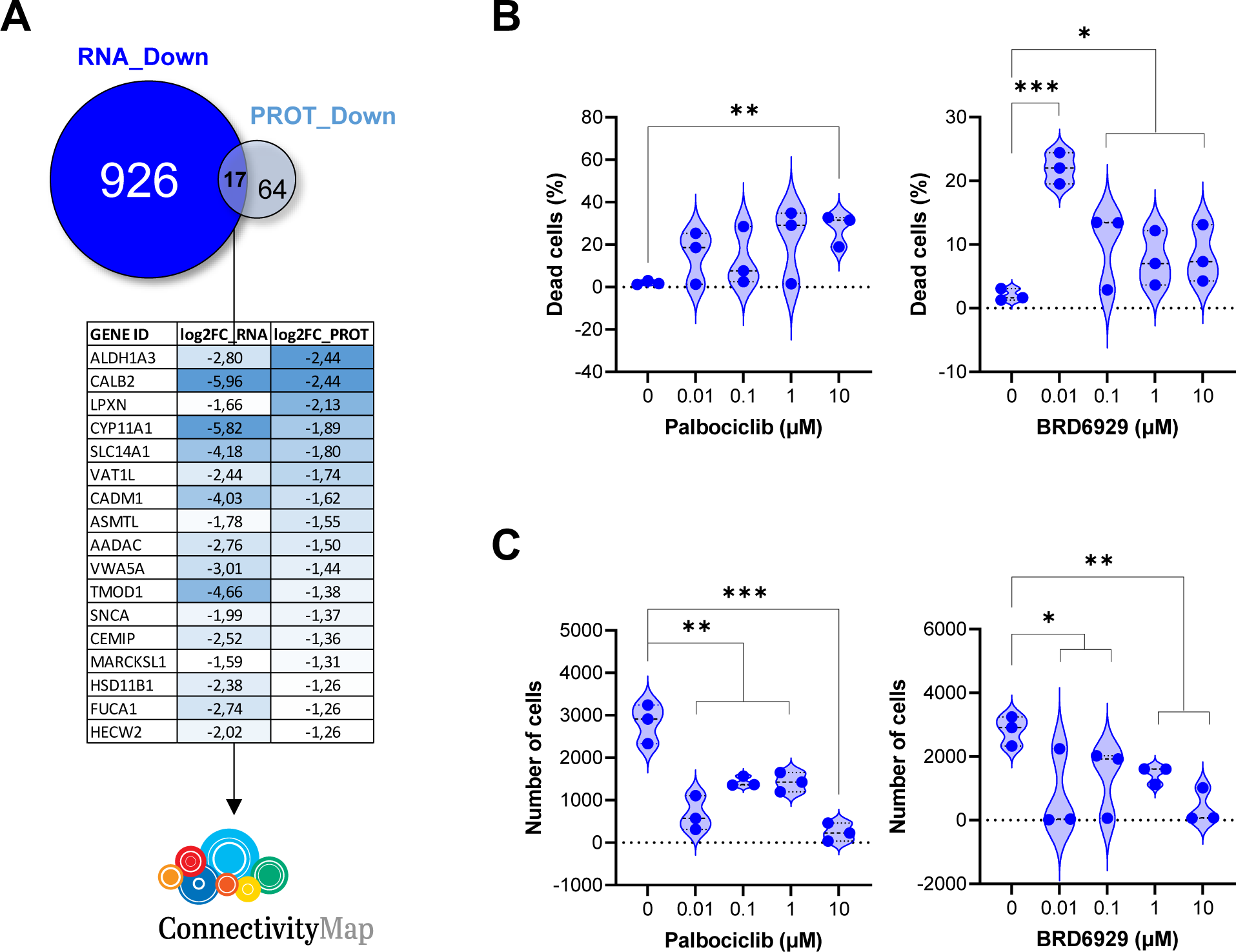
Palbociclib and BRD6929 are two compounds identified with ConnectivityMap that recapitulate in KGN cells the antiproliferative effects induced by the elimination of FOXL2-C134W mutation. (A) Genes differentially downregulated in both the transcriptomic and proteomic analyses when comparing PARENTAL and CRISPR clones. Table shows the fold change downregulation of the 17 common genes. (B) Death cell induced by different concentrations of Palbociclib and BRD6929 in PARENTAL cells 24h post treatment. (C) A significant downregulation of cell number is detected in PARENTAL cells 48 hours after they were treated with Palbociclib and BRD6929.

In summary, the proteomic analysis of CRISPR and PARENTAL clones has allowed us to improve the results obtained at the transcriptomic level. The filtered results have produced a more detailed characterization of the PARENTAL clones and conducted us to find compounds, whose action partially mimic the elimination of the FOXL2-C134W mutation in KGN cells.

## Discussion

### Using CRISPR to study the role of FOXL2-C134W in granulosa cell tumors

The pathognomonic mutation c.402C>G is found in the FOXL2 gene in over 95% of granulosa cell tumors [13]. Since only a few other mutations have been reported in this tumor type [58], they can be considered a highly unique cancer primarily associated with a single mutation in a single gene crucial for the cells of origin, the granulosa cells. Recently, the causal relationship between FOXL2-C134W and AGCTs has been demonstrated through the generation and characterization of a mouse model carrying the equivalent mutation in mice, Foxl2C130W [47]. All female mice carrying this mutation develop ovarian cancers that closely resemble AGCT pathology. We have reached similar conclusions by concurrently developing an equivalent mouse model (Amarilla-Quintana et al., unpublished results).

In this study, we utilized CRISPR technology to target and eliminate the FOXL2 c.402C>G p.C134W mutation in KGN cells. By using mutation-specific guides, we demonstrated that the lack of FOXL2-C134W expression leads to a reduction in the malignant properties of these AGCT cells, indicating their partial dependence on the expression of this variant. This is not the first time CRISPR technology has been employed to target FOXL2 in granulosa tumor cells. Tang and colleagues generated a FOXL2 null clone using CRISPR-mediated techniques [59]. In this clone, abnormal expression of two FOXL2 targets (StAR and CYP19A) was observed, along with decreased cell proliferation. In our study, we generated and analyzed ten clones in which the FOXL2 c.402C>G mutant allele was eliminated using Cas9 complexes, while the wild-type allele remained mostly unaffected by CRISPR activity. The resulting clones behaved similarly to the clone reported by Tang and colleagues, with no major changes observed in apoptosis levels, although significant proliferation inhibition was induced. Additionally, our analysis revealed increased senescence in CRISPR clones compared to control clones. Furthermore, these KGN cells in which the FOXL2 C134W mutation was disrupted exhibited limited invasion capacity both in vitro and in vivo (see Figure 4).

In a more recent study, Heman and colleagues [60] utilized CRISPR technology to revert the mutant allele to the wildtype one. The two wild-type, edited clones they obtained were extensively characterized at the transcriptomic level. Interestingly, they compared their signature profiles with those from FOXL2 null backgrounds and concluded that the FOXL2+/C134W abnormalities are consistent with a gain-of-function scenario.

### An expression signature associated with the FOXL2-C134W mutation

The expression studies conducted in this work have enabled us to characterize the expression signature associated with the presence of the FOXL2-C134W mutation in KGN cells. Transcriptional comparison of PARENTAL (non-edited) and CRISPR (edited) KGN clones indicates that the mutation is primarily associated with the upregulation of cell cycle division genes and the downregulation of estrogen regulators, consistent with a higher proliferative status and a lack of functions related to wild-type, differentiated granulosa cells. Similar transcriptomic analyses have been conducted by others using different approaches. Rosario and colleagues, after overexpressing wild-type and mutant FOXL2 in COV434 cells and silencing mutant FOXL2 expression in KGN cells, demonstrated that many genes regulated by mutant FOXL2 are involved in cell death, proliferation, tumorigenesis, and TGF-β signaling [46]. Years later, Weas-Banks and colleagues reported a transcriptional signature associated with FOXL2-C134W overexpression in human granulosa cells, characterized by epithelial-to-mesenchymal transition and induction of various cytokines, oncogenes, and factors involved in stemness [26]. More recently, Herman and colleagues analyzed the transcriptional consequences of reverting the FOXL2-C134W mutation to wild type and described dysregulated pathways, highlighting the TGF-beta pathway, cell adhesion, and migration processes as the most significantly disrupted [60]. Although there is no major conservation between the differentially expressed genes (DEGs) of all these studies and ours, the enriched terms and pathways are common among them. Indeed, our study also detected enrichment of TGF-beta, stemness, cell adhesion, extracellular matrix, epithelial-to-mesenchymal transition, apoptosis, and migration among the significant outcomes obtained from Oncosignature, Terms, Pathways, or Perturbations sets (Supplementary Tables 4 and 6).

To our knowledge, this is the first study in which transcriptomic and proteomic data are simultaneously obtained and analyzed for granulosa tumor cells. Fewer differentially expressed proteins (DEPs) than DEGs were found in this work. This could be due to the lower power of detection associated with proteomic methodology compared to transcriptomics. However, it is also possible that fewer proteins than transcripts are significantly altered due to post-translational mechanisms that further regulate the transcriptional changes induced by the FOXL2-C134W mutant form. Nevertheless, the differences found between transcriptomic and proteomic data suggest that RNA perturbations should be interpreted with caution even in the study of a transcription factor such as FOXL2. On the other hand, the common deregulated genes and Gene Ontology (GO) terms found between transcriptomic and proteomic studies are of particular value. Some of them have not been previously associated with AGCT, thus opening new avenues for understanding and treating the disease. Additionally, the common GO terms could be of special interest from a therapeutic point of view, as ribosome biogenesis appears to be a key and distinct upregulated feature among AGCT cells. Accumulated recent evidence highlights the importance of ribosome biogenesis in cancer [61], suggesting that further studies should be implemented to explore the potential of targeting ribosome biogenesis in AGCT.

### Therapeutic implications: improving existing approaches and finding new compounds for the treatment of AGCT

One of the main questions we initially posed in this study was whether CRISPR could serve as a therapeutic tool against AGCT. Our results strongly suggest so. By disrupting the FOXL2 c.402C>G p.C134W mutation using CRISPR/Cas9, we observed a significant reduction in the malignant properties of KGN cells both *in vitro* and *in vivo*. Moreover, KGN cells in which the FOXL2-C134W mutation was eliminated became more susceptible to drugs previously used against AGCT, such as Dasatinib and Ketoconazole.

Our expression studies have provided valuable insights that could help the design of future therapeutic strategies. Notably, the higher levels of PDL1 (CD274) in KGN cells compared to those in the edited clones (Figure 5c), suggest the potential clinical benefit of immune checkpoint-based therapies. Additionally, it is noteworthy that two HLA proteins (HLA-DRB3 and HLA-DRb5) were among the top-25 most downregulated proteins in KGN cells (see Figure 6d). Although these proteins have not yet been implicated in immunotherapy, they could potentially impact the effectiveness of such treatments in AGCT.

Furthermore, employing an *in silico* approach, we utilized the signature induced upon the elimination of FOXL2-C134W in KGN cells to identify new compounds specifically targeting them. Remarkably, a fraction (16 out of 100) of the compounds identified through this strategy, overlapped with those reported in a previous drug screening conducted by others [34] (Supplementary Table 9), aiming to uncover new therapeutic opportunities. In our study, HDAC inhibitors, topoisomerase inhibitors, and CDK inhibitors emerged as the most promising categories for future therapeutic strategies, either as single agents or in combination with others.

### Conclusions and limitations of the study

The major limitation of this work is the fact that all experiments were conducted using a single cell line, KGN. Unfortunately, this limitation is unavoidable as the KGN cell line is the only available AGCT cell line. To mitigate this limitation, we utilized multiple clones that underwent the CRISPR process, resulting in clones maintaining their original status (PARENTAL) or losing the FOXL2-C134W mutation (CRISPR).

In this study, we have demonstrated that specific disruption of the FOXL2-C134W mutation in a CRISPR-mediated manner reduces the malignant phenotype of AGCT cells and increases their susceptibility to drugs previously used for AGCT treatment. These findings strongly suggest that AGCT cells are dependent on the FOXL2-C134W mutation.

Furthermore, comparison of CRISPR-edited and parental KGN cells has provided valuable insights into the underlying mechanisms involved in AGCT development and progression and propose new therapeutic strategies for their treatment.

## List of Figures/Tables

Figure 1. Edition of FOXL2 c.402C>G mutation in KGN cells.

Figure 2. Specific FOXL2 c.402C>G mutation elimination causes a decrease in cell growth.

Figure 3. Cell survival and growth regulation pathways are affected by the specific elimination of FOXL2 c.402C>G mutation in KGN cells.

Figure 4. Migration capacity and sensitivity to Dasatinib and Ketokonazole is altered in CRISPR clones.

Figure 5. CRISPR-mediated elimination of FOXL2 c.402C>G mutation induces significant changes in the transcriptomic profile of granulosa tumor cells.

Figure 6. Proteomic changes induced by the CRISPR-mediated elimination of FOXL2 c.402C>G mutation.

Figure 7. Palbociclib and BRD6929 are two compounds identified with ConnectivityMap that recapitulate in KGN cells the antiproliferative effects induced by the elimination of FOXL2-C134W mutation.

Table 1. Resources used in this work.

## Supplementary Materials

Supplementary Figure1. Edition of FOXL2 c.402C>G mutation in KGN cells upon transfection of CRISPR/Cas9 complexes.

Supplementary Figure 2. Validation by RT-qPCR of transcriptomic data.

Supplementary Figure 3. Clustering analyses of transcriptomic data from tumoral and non-tumoral granulosa cell clones.

Supplementary Figure 4. Network clustering of proteomic GO terms from differential expressed proteins. Supplementary Table1. Genomic analysis from pools after gene edition.

Supplementary Table 2. Indels characterization of the reads obtained from amplicon Deep sequencing of the pools after gene edition.

Supplementary Table 3. List of clones generated from pools edited with guides sg1.3 and sg1.4. Supplementary Table 4. List of the DEG found in KGN clones when compared with CRISPR ones.

Supplementary Table 5. List of top25 up (red) and top25 down (blue) regulated genes differentially expressed between PARENTAL and CRISPR clones.

Supplementary Table 6. List of the DEPs found in KGN clones when compared with CRISPR ones. Supplementary Table 7. Genes commonly de-regulated in the transcriptomic and proteomic analyses.

Supplementary Table 8. List of top25 up (red) and down (blue) regulated proteins differentially expressed between PARENTAL and CRISPR clones.

Supplementary Table 9. cMap first 100 ranked compounds that mimic the expression signature induced in KGN cells upon the elimination of FOXL2-C134W mutation.

Supplementary Table 10. Types of compounds that mimic the expression signature induced in KGN cells upon the elimination of FOXL2-C134W mutation.

## Author Contributions

S.A.-Q. acquired and analyzed most of the data. P.M., A.R. and A.M. assist with several molecular experiments. A.M.-C. and R.B. performed and analyzed proteomic studies. M.J.B., S.M. and I.C. carried out transcriptomic analyses. B.V.-M., C.E., D.G.D. and I.H. collaborate in molecular assays. P.C.-S and L.S. performed and analyzed the zebrafish studies. D.M. assisted in all the microscopy related experiments. J.G.-D. and I.P.C. developed the study concept and obtained funding. A.M. and I.P.C. supervised the work, interpreted the data and drafted/edited the manuscript. All authors have read and agreed to the published version of the manuscript.

## Data Availability Statement

The data supporting the findings of this study are available within the article and its Supplementary Material. All transcriptomic and proteomic raw data will be made publicly available in public repositories following peer review and publication.

## Acknowledgments

We want to acknowledge the very important role of patients on the rare disease, scientific works. Specially, we want to deeply thank the support we found on Blanca Antuña. We would like to thank also Vanessa Lafarga for her advice and technical support on the ConnectivityMap studies. We acknowledge grade students Irene Hernández and Andrea González for their enthusiastic participation in some phases of this work.

