## Supplementary material for "CRISPR targeting of FOXL2 c.402C>G mutation reduces malignant phenotype in granulosa tumor cells and identifies anti-tumoral compounds": Supp Figs & Tables

### Supplementary Figures

A

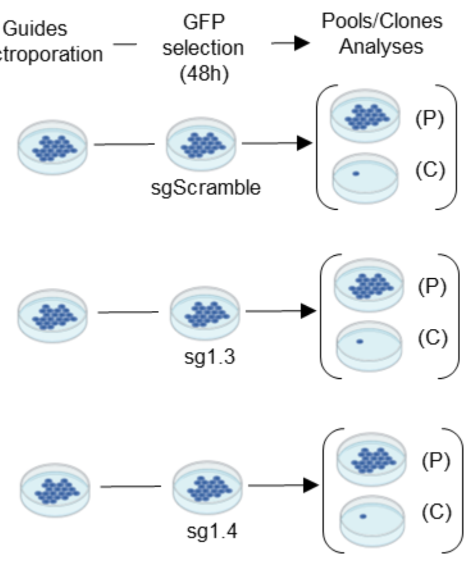


B

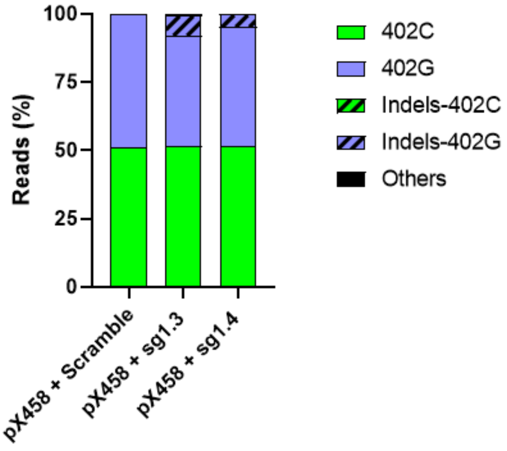


| Reads | px+Scramble | px+sg1.3 | px+sg1.4 |
|---|---|---|---|
| 402 C | 50.92 | 51.57 | 51.45 |
| 402 G | 49.08 | 40.54 | 43.84 |
| Indels in 402 C | 0.00 | 0.00 | 0.00 |
| Indels in 402 G | 0.00 | 7.37 | 4.48 |
| Others | 0.00 | 0.52 | 0.24 |

C

| Indels in 402 G | KGN Cas9 | KGN Cas sg1.3 | KGN Cas sg1.4 | Expected protein |
|---|---|---|---|---|
| c. 399delC | 0 | 75.56 | 0 | p. Trp134Glyfs15* |
| c. 398_399insT | 0 | 7.86 | 0 | p. Trp134Leufs104* |
| c. 398_399insC | 0 | 8.71 | 0 | p. Trp134Leufs104* |
| c. 397_398del GC | 0 | 4.36 | 0 | p. Ala133Leufs104* |
| c. 403_404insA | 0 | 0 | 12.40 | p. Asp136Lysfs242* |
| c. 404delA | 0 | 0 | 55.28 | p. Asp136Thrfs13* |
| c. 404_405delAA | 0 | 0 | 5.28 | p. Glu135Glyfs102* |
| c. 404_406delAAG | 0 | 0 | 27.03 | p. Glu135Aspfs241* |
| c. 403del (>15 bp) | 0 | 3.51 | 0 | |

**Supplementary Figure 1. Edition of FOXL2 c.402C>G mutation in KGN cells upon electroporation of CRISPR/Cas9 complexes. (A)** Timeline for the generation of pools and clones. **(B)** Guide specificity on the FOXL2 c.402C>G mutant allele. Graph represents percentage of reads matching each edited and not edited alleles for the pool of nucleofected cells for each condition (number data represented in bottom table of the same panel). **(C)** Indels identified in nuleofected pools, after activity of each CRISPR/Cas9 complex.

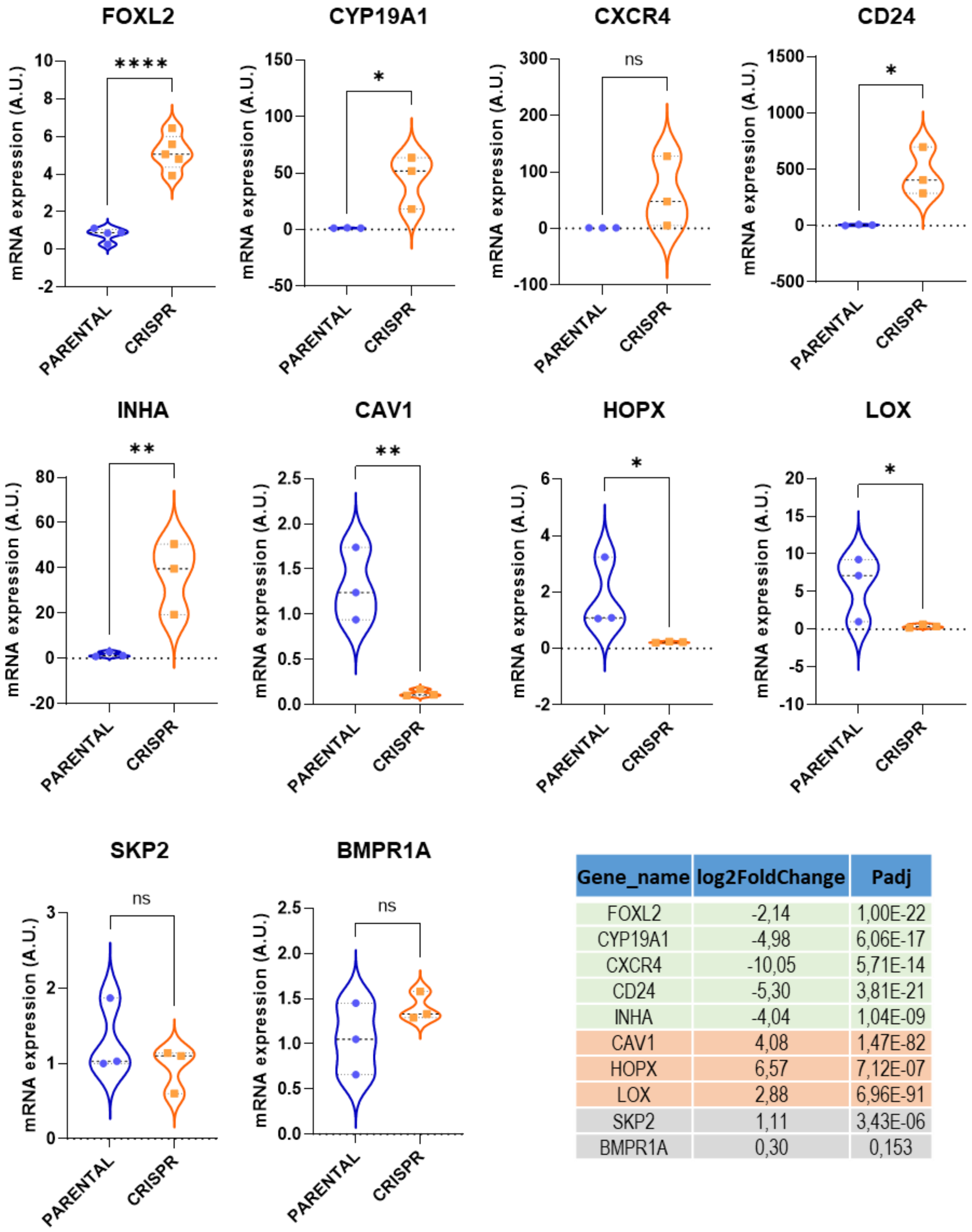


| Gene_name | log2FoldChange | Padj |
|---|---|---|
| FOXL2 | -2,14 | 1,00E-22 |
| CYP19A1 | -4,98 | 6,06E-17 |
| CXCR4 | -10,05 | 5,71E-14 |
| CD24 | -5,30 | 3,81E-21 |
| INHA | -4,04 | 1,04E-09 |
| CAV1 | 4,08 | 1,47E-82 |
| HOPX | 6,57 | 7,12E-07 |
| LOX | 2,88 | 6,96E-91 |
| SKP2 | 1,11 | 3,43E-06 |
| BMPR1A | 0,30 | 0,153 |

**Supplementary Figure 2. Validation by RT-qPCR of transcriptomic data.** Graphs show the results from quantitative PCR results of 10 genes included in the transcriptomic study. FoldChanges and adjusted P values are included in the bottom right table.

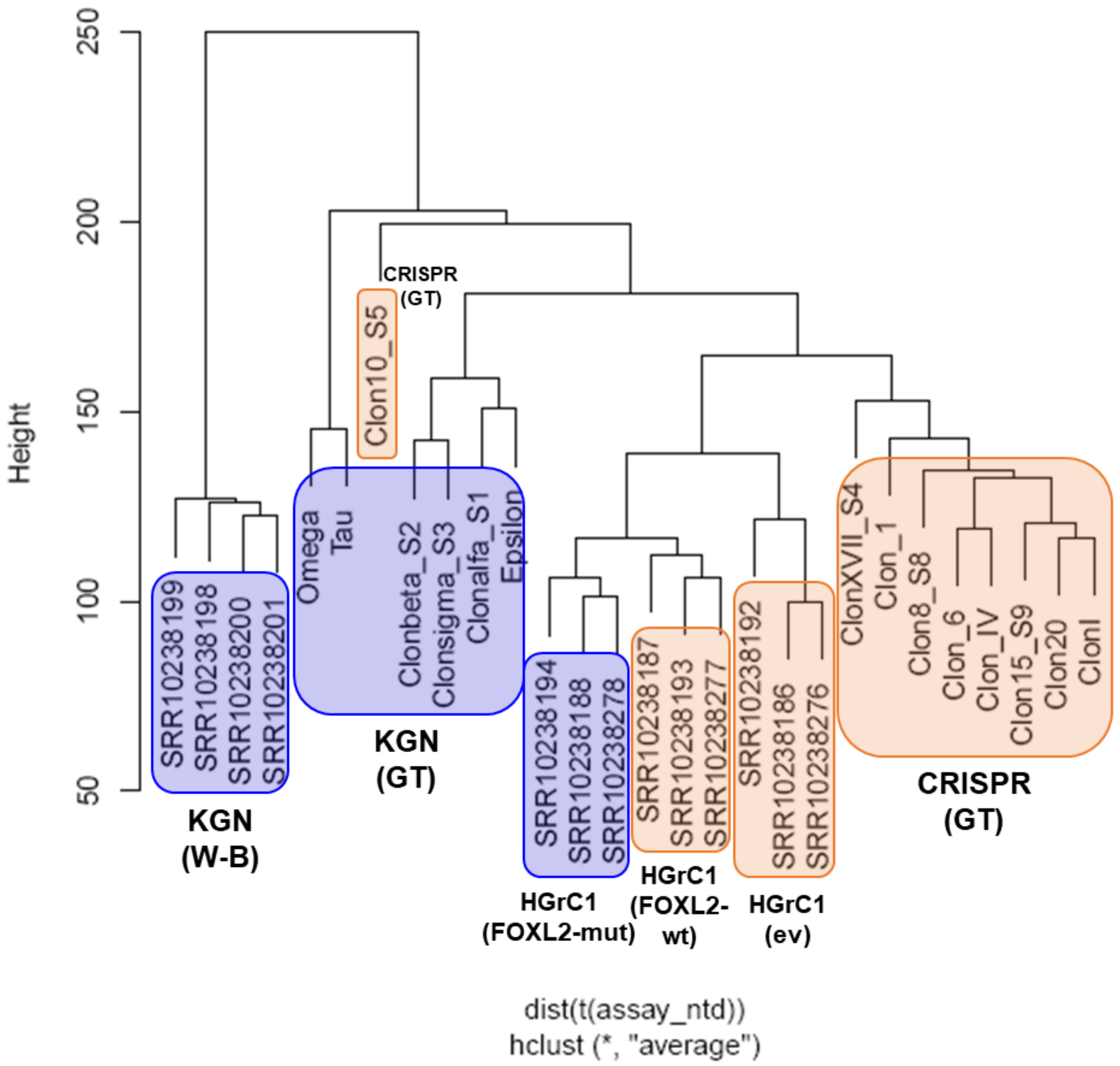


**Supplementary Figure 3. Clustering analyses of transcriptomic data from tumoral and non-tumoral granulosa cell clones.** Parental, KGN(GT), and CRISPR, CRISPR(GT) clones from this study are compared with KGN and HGrC1 granulosa cell lines generated by Weis-Banke et al [56][56]KGN (W-B), empty vector (ev), HGrC1 (FOXL2-mut) and HGrC1 (FOXL2-wt). KGN/Parental genotype cells are marked in blue, and CRISPR/Edited/WT cells are colored in orange.

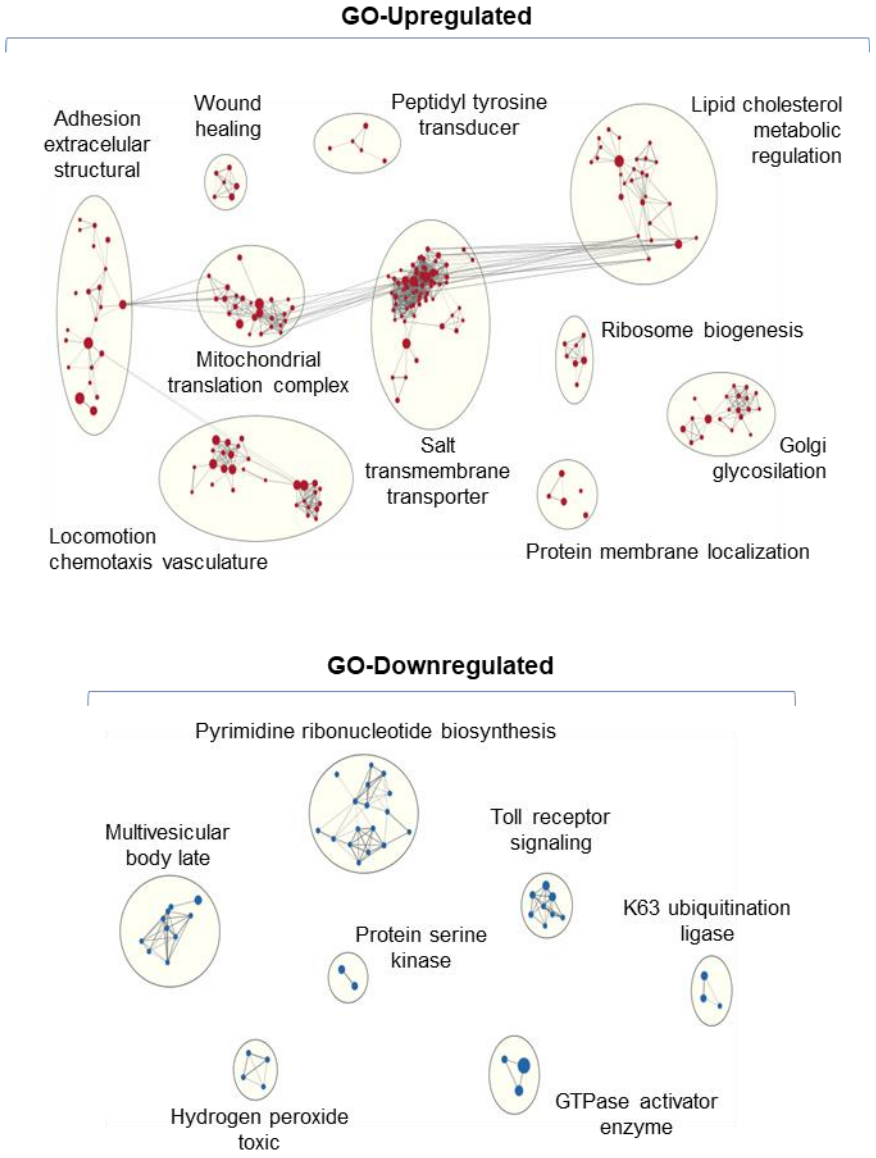


**Supplementary Figure 4. Network clustering of proteomic GO terms. From differentially expressed proteins.** Up and down-regulated, GO terms at q-value<0.05 are shown.

**Supplementary Tables**

**Supplementary Table 1. Genomic analysis from pools after gene edition.** Table includes percentage of reads for the wild-type allele (402C), mutant allele (402G), total editions on mutant (Indels in 402G) and wild-type (Indels in 402C) alleles and, finally, for edited reads in which allele origin cannot be determined (Others).

| **Allele** | % | | |
|---|---|---|---|
| | **KGN Cas9** | **KGN Cas sg1.3** | **KGN Cas sg1.4** |
| **402 C** | 50,21 | 73,85 | 61,24 |
| **402 G** | 49,57 | 8,83 | 3,44 |
| **Indels in 402 C** | 0,00 | 0,00 | 0,00 |
| **Indels in 402 G** | 0,00 | 14,31 | 25,04 |
| **Others** | 0,22 | 3,02 | 10,27 |

**Supplementary Table 2. Indels characterization of the reads obtained from amplicon Deep sequencing of the pools after gene edition.**

| **Indels in 402 G** | % | | | **Expected protein** |
|---|---|---|---|---|
| | **KGN Cas9** | **KGN Cas sg1.3** | **KGN Cas sg1.4** | |
| c. **399delC** | 0 | 65,81 | 1,76 | p. Trp134Glyfs15* |
| c. **392_402del** | 0 | 10,60 | 0 | p. Asp131Glyfs103* |
| c. **398_399insT** | 0 | 7,64 | 0 | p. Trp134Leufs104* |
| c. **398_399insC** | 0 | 3,56 | 0 | p. Trp134Leufs104* |
| c. **399_400delCT** | 0 | 2,57 | 0 | p. Trp134Glyfs103* |
| c. **398_399insA** | 0 | 2,49 | 0 | p. Trp134Leufs104* |
| c. **397_398del GC** | 0 | 2,06 | 0 | p. Ala133Leufs104* |
| c. **398_399insG** | 0 | 1,84 | 0 | p. Trp134Leufs104* |
| c. **395_399del** | 0 | 1,33 | 0 | p. Pro132Leufs104* |
| c. **400delT** | 0 | 1,07 | 0 | p. Trp134Glyfs15* |
| c. **399_401delCTG** | 0 | 1,03 | 0 | p. Trp134Glufs142* |
| **c. 403_404insA** | 0 | 0 | 46,21 | p. Asp136Lysfs242* |
| **c. 404delA** | 0 | 0 | 26,04 | p. Asp136Thrfs13* |
| **c. 404_405delAA** | 0 | 0 | 9,79 | p. Glu135Glyfs102* |
| **c. 404_406delAAG** | 0 | 0 | 6,20 | p. Glu135Aspfs241* |
| **c. 403_404insAA** | 0 | 0 | 5,91 | p. Asp136Lysfs242* |
| **c. 403_404insC** | 0 | 0 | 2,26 | p. Glu135Alafs103* |
| **c. 404_407del** | 0 | 0 | 1,83 | p. Glu135Alafs13* |

**Supplementary Table 3. List of clones generated from pools edited with guides sg1.3 and sg1.4.**

| ID | TYPE | Indel at allele | Genotype | Protein (expected for the edited allele) |
|---|---|---|---|---|
| **K_E1_sg13_3pulso** | CRISPR | 402G | **WT / c.398_399insT** | p. Trp134Leufs104* |
| **K_E1_sg13_1** | CRISPR | 402G | **WT / c. 399delC** | p. Trp134Glyfs15* |
| **K_E1_sg13_6** | CRISPR | 402G | **WT / c. 399delC** | p. Trp134Glyfs15* |
| **K_E1_sg13_8** | CRISPR | **Other** | **WT / c.[397_398GC>TT];[399_402del]; [404_406AAG>CCA]** | p. Ala133Leufs15* |
| **K_E1_sg13_10** | CRISPR | 402G | **WT / *** | p. 376* |
| **K_E1_sg13_11** | CRISPR | 402G | **WT / c. 399delC** | p. Trp134Glyfs15* |
| **K_E1_sg13_12** | CRISPR | 402G | **WT / c. 389_398del** | p. Asp131Glyfs103* |
| **K_E1_sg13_13** | CRISPR | 402G | **WT / c.398_399insT** | p. Trp134Leufs104* |
| **K_E1_sg13_15** | CRISPR | 402G | **WT / c. 399delC** | p. Trp134Glyfs15* |
| **K_E1_sg13_17** | CRISPR | 402G | **WT / c. [399_400insT]; [398C>T]** | p. Trp134Leufs104* |
| **K_E1_sg13_18** | CRISPR | 402G | **WT / c. 389_398del** | p. Asp131Glyfs103* |
| **K_E1_sg13_20** | CRISPR | 402G | **WT / c. 399delC** | p. Trp134Glyfs15* |
| **K_E1_sg14_IIpulso** | CRISPR | 402G | **WT / c.403delG** | p. Glu135Lysfs14* |
| **K_E1_sg14_I** | CRISPR | 402G | **WT / c.405delA** | p. Asp136Thrfs14* |
| **K_E1_sg14_III** | CRISPR | 402G | **WT / c.406_407insAA** | p. Asp136Glufs14* |
| **K_E1_sg14_IV** | CRISPR | **Other** | **WT / c.396_403del** | p. Ala133Argfs102* |
| **K_E1_sg14_V** | CRISPR | 402G | **WT / *** | p. 376* |
| **K_E1_sg14_VII** | CRISPR | 402G | **WT / c. 403_404insA** | p. Asp136Argfs102* |
| **K_E1_sg14_XIV** | CRISPR | 402G | **WT / c. 403_404insA** | p. Asp136Argfs102* |
| **K_E1_sg14_XVI** | CRISPR | 402G | **WT / c. 403_404insA** | p. Asp136Argfs102* |
| **K_E1_sg14_XVII** | CRISPR | 402G | **WT / *** | p. 376* |
| **K_E1_sg14_XVIII** | CRISPR | 402G | **WT / c.406_407insAA** | p. Asp136Glufs14* |
| **K_E1_sg14_XX** | CRISPR | **Other** | **WT / c. 402_404del** | p. *134 |
| **K_E1_sg14_IIIp** | KO | 402C | **c. 404delA** | p. Asp136Thrfs13* |
| **K_E1_sg14_XXI** | KO | 402C | **c. 403_404insA** | p. Asp136Argfs102* |
| **K_E1_sg14_IX** | CRISPR | 402G | **WT / *** | p. 376* |
| **K_E1_sg14_VI** | CRISPR | 402G | **WT / c.396_403del** | p. Ala133Argfs102* |

**Supplementary Table 4. List of the DEG found in KGN clones when compared with CRISPR ones (submitted upon request).**

**Supplementary Table 5. List of top25 up (red) and top25 down (blue) regulated genes differentially expressed between PARENTAL and CRISPR clones.**

| GENE | Function and relationship with cancer | Relation with granulosa cells and/or FOXL2 |
|---|---|---|
| **AOX1** Aldehyde oxidase 1 | It produces hydrogen peroxide and it is implicated in ROS. It has been related with different cancer types including pancreatic cancer (PDAC), clear cell renal cell carcinoma (ccRCC) and colorectal cancer (CRC). | Enriched in heat-stressed granulosa cells (doi: 10.20944/preprints202204.0248.v1) |
| **IFRD1** Interferon Related Developmental Regulator 1 | Protein related to interferon gamma. Transcriptional co-activator/repressor that controls the growth and differentiation of specific cell types during embryonic development and tissue regeneration. Protein expression in colon cancer is associated with poorer patient prognosis. | Ifrd1 mRNA is primarily induced in granulosa cells during the periovulatory period in the rat ovary (doi: 10.1002/MRD.22673). |
| **CAV1** Caveolin 1 | Main component of the caveolae plasma membranes. Negative regulator of RAS signaling. Links integrin to Ras-ERK signaling. It has been related to various cancer types, including head and neck squamous cell carcinoma, melanoma, and Ewing's sarcoma family tumors. | It plays a role in folliculogenesis and female reproduction (doi: 10.1093/HUMREP/DEY299). In bovine ovaries, CAV1 is expressed in granulosa and theca cells of the follicle and large and small cells of the corpus luteum (doi: 10.1002/MRD.20513). |
| **TBC1D9** TBC1 Domain Family Member 9 | Activation of GTPase activity and intracellular protein transport Associated to the inhibition of migratory and invasive capabilities of colorectal cancer cells. Downregulated in breast cancer. | None |
| **CAV2** Caveolin 2 | Major component of the inner surface of caveolae, small invaginations of the plasma membrane, and is involved in essential cellular functions, including signal transduction, lipid metabolism, cellular growth control and apoptosis CAV2 plays a significant role in cancer progression and metastasis in various types of cancer, including head and neck squamous cell carcinoma (HNSCC), breast cancer, and pancreatic cancer. | None |
| **SSX2IP** SSX Family Member 2 Interacting Protein | It is involved in the formation of the actin cytoskeleton at adherens junctions. It is also a centrosome maturation and maintenance factor.<br>Associated with different cancer types including nasopharyngeal carcinoma, acute lymphoblastic leukemia, hepatocellular carcinoma and myeloid leukemia. | None |

| | | |
|---|---|---|
| **ACTA2**<br>Actin Alpha 2, Smooth Muscle | Implicated in actin polymerization and nucleotide binding. Six actin different proteins.<br>High expression of ACTA2 is associated with a worse overall survival and response to immune checkpoint inhibitors in gastric cancer. Also related to relapse in glioma patients, being involved in a higher migration and malignant phenotype. | Upregulated in theca interna when compared with granulosa cells (doi: 10.1371/journal.pone.0119800) |
| **TMEM178B**<br>Transmembrane Protein 178B | This transmembrane protein is a negative regulator of inflammatory cytokine production.<br>A TMEM178B-BRAF fusion has been reported in a malignant melanoma. | None |
| **RAB3B**<br>RAB3B, Member RAS Oncogene Family | Enables GDP binding activity; GTPase activity; and myosin V binding activity. Involved in several processes, including positive regulation of dopamine uptake involved in synaptic transmission; regulation of synaptic vesicle cycle; and regulation of vesicle size.<br>Up-regulated in different cancer types including gastric cancer, hepatoma cells and gliomas. | None |
| **MYRF**<br>Myelin Regulatory Factor | Transcription factor involved in olygodendrocyte differentiation, and regulator of Cardiac and Early Gonadal Development.<br>Related to pancreatic cancer progression and testicular germ cell tumors. | None |
| **LOX**<br>ysyl Oxidase | Tumor suppresor via canonical and non-canonical TGFB signaling.<br>It promotes the crosslinking between ECM collagen type I, collagen type III and elastin.<br>Involved in multple cancers. | It plays a critical role in the regulation of granulosa cell differentiation being involved in tissue remodeling and ECM (doi: 10.1530/REP-16-0254) |
| **VSNL1**<br>Visinin Like 1 | Neuronal calcium sensor protein. Directly or indirectly regulates the activity of adenylyl cyclase.<br>Associated with cholangiocarcinoma, colorectal carcinogenesis, glioblastoma and squamous cell carcinoma. | None |
| **DNAH11**<br>Dynein Axonemal Heavy Chain 11 | Microtubule-dependent motor ATPase involved in the movement of respiratory cilia.<br>It has been connected to colon cancer, pituitary adenomas, ovarian and breast cancer. | None |
| **AXL**<br>AXL Receptor Tyrosine Kinase | Tyr-K receptor. Involved in several cellular functions including growth, migration, aggregation and anti-inflammation in multiple cell types.<br>Associated with different tumor types. | Involved in the viability of hypoxic granulosa cells (doi: 10.1002/jcp.31162). Expressed by granulosa cells at high levels throughout antral follicle development. |

| | | |
|---|---|---|
| **CARMN** Cardiac Mesoderm Enhancer-Associated Non-Coding RNA | Predicted to be involved in regulation of gene expression. Associated with multiple tumor types. | None |
| **CSPG4** Chondroitin Sulfate Proteoglycan 4 | CSPG4 can mediate intracellular signalling downstream of growth factor receptor and integrin interactions, potentiating communication between the extracellular and intracellular compartments of the cell. Related to several tumor types. | Upregulated in TGFβ-induced GCTs, when compared with wild type granulosa cells (doi: 10.3390/cancers14092184) |
| **TAGLN** Transgelin | It is involved in calcium-independent smooth muscle contraction. It acts as a tumor suppressor, and the loss of its expression is an early event in cell transformation and the development of some tumors, coinciding with cellular plasticity. | Upregulated in TGFβ-induced GCTs when compared with wild type granulosa cells (doi: 10.3390/cancers14092184). |
| **LCP1** Lymphocyte Cytosolic Protein 1 | Actin-binding protein. It plays a role in the activation of T-cells in response to costimulation through TCR/CD3 and CD2 or CD28 and modulates the cell surface expression of IL2RA/CD25 and CD69. Associated with cancer. | None |
| **CD274** PD-L1 | PDL-1. Inhibitory receptor ligand that is expressed by hematopoietic and non-hematopoietic cells, such as T cells and B cells and various types of tumor cells. Key immune checkpoint player considered to be prognostic in many types of human malignancies, including colon cancer and renal cell carcinoma. | Not expressed in granulosa cell baseline tumors (doi: 10.1007/s10637-020-01043-9) |
| **JPH2** Junctophilin 2 | Component of junctional complexes that mediate cross talk between cell surface and intracellular ion channels. Abnormally expressed in leimyosarcoma and spcifically methylated in gastric cancer. | Upregulated in TGFβ-induced GCTs when compared with wild type granulosa cells (doi: 10.3390/cancers14092184). |
| **FGF1** Fibroblast Growth Factor 1 | Fibroblast growth factor 1. Mitogenic and cell survival activities Correlates with poor survival in various cancers and resistance to platinum-based chemotherapy of serous cancers. | Favors survival of COV434 granulosa tumor cells upon etoposide treatment (doi: 10.1038/s41389-018-0033-y). Involved in the differentiation of ovarian follicles. |
| **FBN2** Fibrillin 2 | Component of connective tissue microfibrils that may be involved in elastic fiber assembly. Abnormal methylation/expression in diferent tumor types. | None |
| **TRPC4** Transient Receptor Potential Cation Channel Subfamily C Member 4 | It forms a non-selective calcium-permeable cation channel that is activated by Gq-coupled receptors and tyrosine kinases and plays a role in multiple processes including endothelial permeability, vasodilation, neurotransmitter release and cell proliferation. Associated with prognosis, tumor microenvironment and treatment sensitivity. | None |

| | | |
|---|---|---|
| **SLC8A1**<br>Solute Carrier Family 8 Member A1 | Low affinity, high capacitance calcium antiporter membrane protein that functions to regulate intracellular calcium concentrations. Related to Megacolon and Penile cancer. | Target gene of FOXL2 (PANTHER CLASS as: cell communication and signal transduction) (doi: 10.7554/eLife.04207) |
| **LMOD1**<br>Leiomodin-1 | Required for proper contractility of visceral smooth muscle cells, it mediates nucleation of actin filaments.<br>Deregulated in endometrial cancer (doi: 10.1038/s41598-020-66872-3). | Involved in ovulation in mouse ovary (doi: 10.1101/2023.08.21.554210). |
| **SMIM3**<br>Small integral membrane protein 3 | Transmembrane protein involved in cell channel regulation and associated with neuronal differentiation.<br>Related to pheochromocytonas, oral squamous cell carcinomas and aucte myeloid leukemia. | None |
| **KSR1**<br>Kinase suppresor of Ras1 | Enables 14-3-3 protein binding activity, ATP binding activity, and protein C-terminus binding activity. Involved in positive regulation of MAPK cascade.<br>Associated with different tumor types. | None |
| **AHRR**<br>Aryl Hydrocarbon Receptor Repressor | It participates in the aryl hydrocarbon receptor (AhR) signaling cascade, which mediates dioxin toxicity, and is involved in regulation of cell growth and differentiation.<br>Related to cancer at different levels. | Activated in granulosa cells upon exposure to contaminants (doi: 10.1016/j.tox.2017.07.003) |
| **HTR7**<br>5-Hydroxytryptamine Receptor 7 | Serotonin receptor.<br>Abnormaly regulated in breast, gastric, laryngeal cancers and acute myeloid leukemia. | None |
| **LIMCH1**<br>LIM And Calponin Homology Domains 1 | Enables myosin II head/neck binding activity. Involved in several processes, including cytoplasmic actin-based contraction involved in cell motility; positive regulation of stress fiber assembly; and regulation of focal adhesion assembly.<br>Related with some cancer types. | None |
| **ARL4C**<br>ADP Ribosylation Factor Like GTPase 4C | Member of the ADP-ribosylation factor family of GTP-binding proteins.<br>Related with cancer processes. | None |
| **GPRC5C**<br>G Protein-Coupled Receptor Class C Group 5 Member C | It may mediate the cellular effects of retinoic acid on the G protein signal transduction cascade.<br>Some roles reported in cancer. | Upregulated in KGN treated with forskolin (doi: 10.1210/en.2004-0889) |
| **UST**<br>Uronyl 2-Sulfotransferase | It transfers sulfate to the 2-position of uronyl residues, such as iduronyl residues in dermatan sulfate and glucuronyl residues in chondroitin sulfate.<br>It may play a role in cancer progression and could potentially serve as a biomarker for predicting patient survival in certain types of cancer. | None |

| | | |
|---|---|---|
| **ABCC4**<br>ATP Binding Cassette Subfamily C Member 4 | It transports various molecules across extra- and intra-cellular membranes.<br>It is involved in muti-drug resistance.<br>It plays an important role in cancer progression and may serve as a potential therapeutic target in certain types of cancer. | Mediator of ovulation in granulosa cells. Involved in prostaglandin signaling (doi: 10.1096/fj.202101931RR) |
| **CDH12**<br>Cadherin 12 | A type II classical cadherin of the cadherin superfamily. It mediates calcium-dependent cell-cell adhesion and appears to be expressed specifically in the brain.<br>It appears to be involved in cancer progression and may serve as a potential therapeutic target in these malignancies. | Granulosa cell marker (doi:10.3389/fendo.2019.00832) |
| **CYP11A1**<br>Cytochrome P450 Family 11 Subfamily A Member 1 | Member of cytochrome P450 superfamily of enzymes, which catalyze many reactions involved in drug metabolism and synthesis of cholesterol, steroids and other lipids. This protein localizes to the mitochondrial inner membrane and catalyzes the conversion of cholesterol to pregnenolone, the first and rate-limiting step in the synthesis of the steroid hormones.<br>It has been implicated in different cancers (breast, kidney, renal, squamous cell and skin). | It plays a crucial role in steroid hormone synthesis in granulosa cells. Increase expression in GCs undergoing luteinization in ovulation (doi: 10.1210/en.2016-1264. Epub 2016 Jul 18). No expression in granulosa cell tumors (doi: 10.1158/0008-5472.CAN-05-1024). FOXL2 represses the activity of the mouse Cyp11a1 promoter (doi: 10.1530/REP-11-0259). |
| **STC2**<br>Stanniocalcin 2 | Secreted, homodimeric glycoprotein that is expressed in a wide variety of tissues and may have autocrine or paracrine functions. Upregulated in various cancers, it is a biomarker for many of them. | Involved in the regulation of apoptosis and autophagy in granulosa cells (doi: 10.1016/j.yexcr.2023.113473) |
| **TSKU**<br>Tsukushi, Small Leucine Rich Proteoglycan | It enables transforming growth factor beta binding activity. Involved in organ development and cholesterol efflux and homeostasis.<br>TSKU expression is associated with poor overall survival in non-small cell lung cancer. | None |
| **GNG2**<br>G Protein Subunit Gamma 2 | One of the gamma subunits of a guanine nucleotide-binding protein. Involved in signaling mechanisms across membranes.<br>It acts as a tumor suppresor in breast cancers (doi: 10.1038/s41419-022-04690-3). Low expression levels in malignant melanomas. | Upregulated in granulosa cells from primary follicles (doi:10.3389/fcell.2018.00085) |
| **COL4A5**<br>Collagen Type IV Alpha 5 Chain | One of the six subunits of type IV collagen, the major structural component of basement membrane.<br>Related to progression of lung and breast cancer. | None |
| **CSGALNACT1**<br>Chondroitin Sulfate N-Acetylgalactosaminyltransferase 1 | It transfers N-acetylglucosamine (GalNAc) to the core tetrasaccharide linker and to elongating chondroitin sulfate chains in proteoglycans.<br>Few works asscociate this gene with cancer. | Involved in maturation of follicular granulosa cells in buffalo (doi:10.1186/s12864-018-5208-6) |

| | | |
|---|---|---|
| **NTNG2**<br>Netrin G2 | Predicted to be involved in several processes, including basement membrane assembly; cell morphogenesis involved in differentiation; and regulation of cell projection organization. Moderate association with different cancer types. | None |
| **NCK2**<br>NCK Adaptor Protein 2 | Member of NCK family of adatpor proteins. Bind and recruits various proteins involved in regulation of receptor protein tyr-k. Few reports associate this gene with cancer. | None |
| **SLC14A1**<br>Solute Carrier Family 14 Member 1 | Membrane transporter that mediates urea transport in erythrocytes. Biomarker in some cancers. Associated with the progression of different tumors. | None |
| **JCAD**<br>Junctional Cadherin 5 Associated | Endothelial cell to cell junction protein.<br>It plays a role in the development and progression of certain types of cancer. | None |
| **CD55**<br>CD55 Molecule | Glycoprotein involved in the regulation of the complement cascade. Weakly associated with cancer. | Recently identified as a cell surface marker that can be used to isolate early-stage granulosa cells (doi: 10.1111/cpr.13589) |
| **TMOD1**<br>Tropomodulin 1 | Actin-capping protein that regulates tropomyosin by binding to its N-terminus, inhibiting depolymerization and elongation of the pointed end of actin filaments and thereby influencing the structure of the erythrocyte membrane skeleton.<br>It may play a role in promoting cancer cell growth and could be a potential target for therapeutic interventions. | Involved in Regulation of Ovarian Granulosa Cell Morphogenesis, Development and Differentiation (doi: 10.3390/ijms20163966) |
| **CALB2**<br>Calbindin 2 | Intracellular calcium-binding protein belonging to the troponin C superfamily. This protein plays a role in diverse cellular functions, including message targeting and intracellular calcium buffering. It plays a role in cancer progression and may serve as a potential biomarker or therapeutic target in certain cancer types. | Useful marker for GCTs (doi: 10.1309/GRH4-JWX6-J9J7-QQTA) |
| **AC022075.1** | Autophagy-related lncRNAs.<br>Associated with neuroblastoma. | None |
| **BMF**<br>Bcl2 Modifying Factor | It has been shown to bind BCL2 proteins and function as an apoptotic activator. This protein is found to be sequestered to myosin V motors by its association with dynein light chain 2, which may be important for sensing intracellular damage and triggering apoptosis.<br>Downregulation associate with progression of some cancer types. | Associated with follicular development (doi: 10.3390/ijms24010401) |

**Supplementary Table 6. List of the DEPs found in KGN clones when compared with CRISPR ones (submitted upon request).**

**Supplementary Table 7. Genes commonly de-regulated in the transcriptomic and proteomic analyses.** Up-regulated genes are in red and down-regulated genes in blue. The list of genes is ordered according to log2foldchange in the proteomic study.

| Gene | log2Foldchange RNA | log2Foldchange PROT |
|---|---|---|
| RARRES2 | 8,84 | 5,97 |
| FABP3 | 3,92 | 4,50 |
| LMOD1 | 4,65 | 4,12 |
| COL1A2 | 5,61 | 3,70 |
| CSPG4 | 6,94 | 3,25 |
| CAV1 | 4,08 | 3,24 |
| ALDH1L2 | 1,64 | 3,14 |
| TAGLN | 6,08 | 3,04 |
| PTN | 1,77 | 2,89 |
| TNC | 2,94 | 2,89 |
| PODXL | 4,38 | 2,74 |
| FLNC | 4,64 | 2,50 |
| CYSTM1 | 1,54 | 2,50 |
| COL1A1 | 2,20 | 2,49 |
| NCALD | 2,31 | 2,47 |
| CAV2 | 2,45 | 2,44 |
| TPST2 | 1,75 | 2,32 |
| ALDH1B1 | 2,60 | 2,23 |
| LCP1 | 4,61 | 2,13 |
| ENAM | 3,90 | 2,07 |
| SORBS1 | 4,86 | 2,07 |
| GAS6 | 1,50 | 2,07 |
| HSPA2 | 2,20 | 2,01 |
| JPH2 | 3,63 | 1,95 |
| ALCAM | 1,60 | 1,92 |
| FAS | 1,62 | 1,82 |
| MYOF | 2,09 | 1,81 |
| JAG1 | 2,51 | 1,72 |
| COL4A1 | 1,82 | 1,72 |
| GPC4 | 2,56 | 1,64 |
| GPC1 | 1,53 | 1,64 |
| CPA4 | 3,23 | 1,57 |
| ITGA5 | 1,70 | 1,55 |
| MYL9 | 3,41 | 1,52 |
| NT5E | 2,45 | 1,51 |
| PTGIS | 2,69 | 1,51 |
| ASMTL | -1,78 | -1,55 |
| CADM1 | -4,03 | -1,62 |
| VAT1L | -2,44 | -1,74 |
| SLC14A1 | -4,18 | -1,80 |
| CYP11A1 | -5,82 | -1,89 |
| LPXN | -1,66 | -2,13 |
| CALB2 | -5,96 | -2,44 |
| ALDH1A3 | -2,80 | -2,44 |

**Supplementary Table 8. List of top25 up (red) and down (blue) regulated proteins differentially expressed between PARENTAL and CRISPR clones.**

| GENE | Function and relationship with cancer | Relation with granulosa cells and/or FOXL2 | Common in Transcriptomic & Proteomic DEG |
|---|---|---|---|
| **RARRES2** Retinoic Acid Receptor Responder 2 | Secreted chemotactic protein that initiates chemotaxis via the ChemR23 G protein-coupled seven-transmembrane domain ligand. This protein is involved in regulation of adipogenesis, energy metabolism, and inflammation.<br>Related to ovarian cancer among others. | Regulates granulosa cells apoptosis during folliculogenesis by upregulating p53 and p21 (doi: 10.1002/mrd.23241). (doi: 10.1095/biolreprod.113.117044) | **Yes** |
| **FABP3**<br>Fatty Acid Binding Protein 3 | Participates in uptake, transport, and metabolism of long-chain fatty-acids in the cells.<br>Tumor suppressor in breast cancer. | Participates in KGN cell proliferation and also increases the expression of aromatase CYP19 (doi: 10.1093/jmcb/mjaa044) | **Yes** |
| **LMOD1** Leiomodin-1 | Required for proper contractility of visceral smooth muscle cells, it mediates nucleation of actin filaments.<br>Deregulated in endometrial cancer (doi: 10.1038/s41598-020-66872-3). | Involved in ovulation in mouse ovary (doi: 10.1101/2023.08.21.554210). | **Yes (top-25)** |
| **COL1A2**<br>Pro-alpha2 chain of type I collagen | Found in connective tissues, bone, cornea, dermis and tendon.<br>Mutations and abnormal expression can cause instability of the extracelular matrix, increasing proliferation and invasion of tumoral cells. | Indicates oocyte quality, being upregulated in granulosa cells (doi: 10.1071/RD14452; 10.1016/j.rbmo.2021.05.018). FoxL2 (wild-type form) affects its transcription and expression, being increased when its knock-down and affecting follicular depletion during reproductive aging. Mutation FoxL2 C134W does not affect expression levels (doi: 10.1016/j.ydbio.2016.05.022) | **Yes** |
| **CSPG4**<br>Chondroitin Sulfate Proteoglycan 4 | It can mediate intracellular signalling downstream of growth factor receptor and integrin interactions, potentiating communication between the extracellular and intracellular compartments of the cell. Related to several tumor types. | Upregulated in TGFβ-induced GCTs, when compared with wild type granulosa cells (doi: 10.3390/cancers14092184) | **Yes (top-25)** |
| **CAV1**<br>Caveolin 1 | Main component of the caveolae plasma membranes. Negative regulator of RAS signaling. Links integrin to Ras-ERK signaling. It has been related to various cancer types, including head and neck squamous cell carcinoma, melanoma, and Ewing's sarcoma family tumors. | It plays a role in folliculogenesis and female reproduction (doi: 10.1093/HUMREP/DEY299). In bovine ovaries, CAV1 is expressed in granulosa and theca cells of the follicle and large and small cells of the corpus luteum (doi: 10.1002/MRD.20513). | **Yes (top-25)** |
| **ALDH1L2** Aldehyde Dehydrogenase 1 Family Member L2 | This mitochondrial enzyme takes part in the folate metabolism (doi: 10.1074/jbc.M110.128843). | Related to cell cycle and apoptosis in primordial follicles (doi: 10.1093/toxsci/kfs137) | **Yes** |

| | | | |
|---|---|---|---|
| **TAGLN** Transgelin | It is involved in calcium-independent smooth muscle contraction. It acts as a tumor suppressor, and the loss of its expression is an early event in cell transformation and the development of some tumors, coinciding with cellular plasticity. | Upregulated in TGFβ-induced GCTs when compared with wild type granulosa cells (doi: 10.3390/cancers14092184). | **Yes (top-25)** |
| **ALDH2** Aldehyde Dehydrogenase 2 or Mitochondrial | Mitochondrial isoform that participates in alcohol metabolism. | Deregulated in granulosa cells from PCOS patients (at protein level) (doi: 10.1177/11795514231206732) | No |
| **PTN** Pleiotrophin | Secreted heparin-binding growth factor that participates in cell growth, migration, angiogenesis ant tumorigenesis. | In the ovary is associated with stromal cell genes with COL1A1 (doi: 10.1038/s41421-022-00492-1). Under FOXL2 knock-down, this gene is upregulated in granulosa cells (doi: 10.1186/1471-213X-9-36) | **Yes** |
| **TNC** Tenascin C | Component of the extracelullar matrix. Contributes to elongation and migration of endothelial cells during angiogenesis in tumors. In ovarian tumors appears to ve overexpressed in the stroma and is also involved in invasion (doi: 10.1371/journal.pone.0136473). | None | **Yes** |
| **MCAM** Melanoma Cell Adhesion Molecule | Involved in glomerular filtration and vascular wound healing. Acts upstream of or within angiogenesis.<br>Biomarker of uveal melanoma. | Protein involved in neovascularization during the formation of corpus luteum in the human ovary, and defined as a FOXL2 target gene (doi: 10.1093/molehr/gag042; 10.7554/elife.04207) | No |
| **CD99** T-Cell Surface Glycoprotein E2 | Cell surface glycoprotein involved in leukocyte migration, transmembrane protein transport, rearrange cytoskeleton, T-cell adhesion and cell death.<br>Also plays an important role in the identification and diagnosis of different ovary tumors. | Granulosa cell tumors are positive for this cell marker, that is characteristic of sex cord-stromal tumors (SCSTs) (PMID: 9720506.) Granulosa cell tumor marker along with activin and inhibin-$\alpha$ (doi: 10.5858/2000-124-0563-IOISIA) | No |
| **PODXL** Podocalyxin | Involved in cell adhesion and morphology in cancer progression by its interactions with actin-binding protein EZR, that increases activation of MAPK/PI3K pathway. | Related to ovarian cancer via activation by a miRNA (doi: 10.1007/s43032-020-00366-5) | **Yes** |
| **SDPR** Serum Deprivation Response Protein (Cavin-2) | Protein increased in serum-starved cells and required for caveola formation. | None | No |
| **CD36** Platelet glycoprotein 4 | Glycoprotein of the platelet surface, serve as a receptor of thrombospondin in various cell lines | Overexpressed in granulosa cells of obese patients, causing malfunction of these cells and problems in fertility and ovarian function (doi: 10.1071/RD18292). Takes part in steroidogenesis and cholesterol pathways in granulosa cells by increasing its expression as a fatty acid transporter (doi: 10.1186/s40104-021- | No |

| | | | |
|---|---|---|---|
| | | 00660-5). Increased expression of CD36 reduces FOXL2 (doi: 10.1186/s12958-017-0276-z) | |
| **CYSTM1** Cysteine-rich and transmembrane domain-containing protein 1 | Implicated in immune system pathways. Biomarker of hepatocellular carcinoma | None | **Yes** |
| **FLNC** Filamin C | Related to myopathies and other muscular phenotypes. | Differentially expressed at transcriptomic level, when compared with *wild-type* FOXL2. It is associated with tumorigenesis due to its interaction with the extracellular matrix (doi: 10.1186/s12967-021-02754-0) | **Yes** |
| **COL1A1** Collagen Type I Alpha 1 Chain | Most abundant protein in the extracellular matrix. | Implicated in follicular phase, accumulating in cells due to its regulation via TGFβ signaling (doi: 10.1096/fj.202001377R). | No |
| **NCALD** Neurocalcin Delta | It is thought to be a regulator of G protein-coupled receptor signal transduction. Associated with ovarian cancer. It's lower expression in this case is related to chemotherapy resistance (doi: 10.1002/jcb.29670; 10.1186/s13048-020-00635-6) | None | **Yes** |
| **CAV2** Caveolin 2 | Major component of the inner surface of caveolae, small invaginations of the plasma membrane, and is involved in essential cellular functions, including signal transduction, lipid metabolism, cellular growth control and apoptosis. It plays a significant role in cancer progression and metastasis in various types of cancer, including head and neck squamous cell carcinoma (HNSCC), breast cancer, and pancreatic cancer. | None | **Yes (top-25)** |
| **TPST2** Tyrosylprotein Sulfotransferase 2 | This type II integral membrane protein is found in the Golgi body catalyzes the O-sulfation of tyrosine residues within acidic regions of proteins. | None | **Yes** |
| **ALDH1B1** Aldehyde dehydrogenases family of proteins | Aldehyde dehydrogenase involved in involved in the metabolism of corticosteroids, biogenic amines, neurotransmitters, and lipid peroxidation. Contributes to proliferation of pancreatic cancer. | Downregulated in cumulus cells (doi: 10.1093/biolre/ioab163) | **Yes** |
| **LCP1** Lymphocyte Cytosolic Protein 1 | Actin-binding protein. It plays a role in the activation of T-cells in response to costimulation through TCR/CD3 and CD2 or CD28 | None | **Yes** |

| | | | |
|---|---|---|---|
| | and modulates the cell surface expression of IL2RA/CD25 and CD69. Associated with cancer. | | |
| **ENAM** Enamelin | Plays a role in the development of theet matrix. Downregulated in renal cancer tissues and also involved in inhibition of proliferation in clear cell renal carcinoma. | None | **Yes** |
| **ALDH1A3** Aldehyde dehydrogenase 1 family member A3 | Catalyzes the NAD-dependent oxidation of aldehyde substrates. Promotes cancer cell growth and invasion inducing gene expression via retinoic acid. | Participates during steroidogenesis in the ovary (10.1016/j.jare.2023.06.002). ALDH1A3 enhances expression of ovarian genes in Sertoli cells, differentiating them into granulosa cells (doi: 10.1016/j.ydbio.2017.02.015) | **Yes** |
| **CALB2** Calbindin 2 **(Calretinin)** | Intracellular calcium-binding protein belonging to the troponin C superfamily. This protein plays a role in diverse cellular functions, including message targeting and intracellular calcium buffering. It plays a role in cancer progression and may serve as a potential biomarker or therapeutic target in certain cancer types. | Useful marker for GCTs (doi: 10.1309/GRH4-JWX6-J9J7-QQTA) | **Yes (top-25)** |
| **TXNIP** Thioredoxin-interacting protein | A thiol-oxidoreductase that is a major regulator of cellular redox signaling which protects cells from oxidative stress. Overexpression of this protein induces cell cycle arrest. | Downregulated in normal granulosa cells and up-regulated in PCOS granulosa patients, generating granulosa cell dysfunction by activation of NLRP3 inflammasome) (doi: 10.1016/j.mce.2022.111824) | No |
| **LPXN** Leupaxin | Focal-adhesion-associated adaptor-protein family member. | Upregulated with CD36 in PCOS patients with obesity (doi: 10.1071/RD18292) | **Yes** |
| **EEF1A2** Elongation factor 1 alpha 2 | Promotes binding of aminoacyl-tRNA to the A-site of ribosomes in protein synthesis. Overexpressed in breast cancer, where is involved in migration and filopodia stimulation in a PI3K-Akt dependent manner. | Critical in the development of ovarian cancer and in Juvenile Granulosa cell tumors, causing cell migration (doi: 10.1016/j.ebiom.2015.03.002) | No |
| **SAT2** Serine acetyltransferase 2 | Catalyzes the N-acetylation of the amino acid thialysine (S-(2-aminoethyl)-L-cysteine) and it is located in exosomes. | None | No |
| **CYP11A1** Cytochrome P450 Family 11 Subfamily A Member 1 | Member of cytochrome P450 superfamily of enzymes, which catalyze many reactions involved in drug metabolism and synthesis of cholesterol, steroids and other lipids. This protein localizes to the mitochondrial inner membrane and catalyzes the conversion of cholesterol to pregnenolone, the first and rate-limiting step in the synthesis of the steroid hormones. It has been implicated in different cancers (breast, kidney, renal, squamous cell and skin). | It plays a crucial role in steroid hormone synthesis in granulosa cells. Increase expression in GCs undergoing luteinization in ovulation (doi: 10.1210/en.2016-1264. Epub 2016 Jul 18). No expression in granulosa cell tumors (doi: 10.1158/0008-5472.CAN-05-1024). FOXL2 represses the activity of the mouse Cyp11a1 promoter (doi: 10.1530/REP-11-0259). | **Yes (top-25)** |

| | | | |
|---|---|---|---|
| **HLA-DRB5; HLA-DRB3** Major Histocompatibility Complex Clas II DR Beta 3 and 5 | HLA class II beta chain paralogues. They play a central role in the immune system by presenting peptides derived from extracellular proteins. Associated with gliomas with aggressive phenotypes. | None | No |
| **SLC14A1** Solute Carrier Family 14 Member 1 | Membrane transporter that mediates urea transport in erythrocytes. Biomarker in some cancers. Associated with the progression of different tumors. | None | **Yes (top-25)** |
| **SERPINB2** Serpin Family B Member 2 | Protein located on the external side of the plasma membrane, with endopeptidase activity, that promotes cell survival and it is associated with cytokine signlaing.<br>Overexpression inhibits apoptosis and promotes cell survival (doi: 10.1096/fj.202001377R). | Expression of this proteins depends on the stage of the follicular cycle. Highly upregulated in pregnant women (doi: 10.1016/j.theriogenology.2020.02.044). FOXL2 is defined as a positive regulator of this gene (doi: 10.3390/cancers11040499) | No |
| **LACC1** Laccase Domain Containing 1 | Oxidoreductase that promotes fatty-acid oxidation, with concomitant inflammasome activation, mitochondrial and NADPH-oxidase-dependent reactive oxygen species production. | None | No |
| **VAT1L** Vesicle amine transport 1 | Oxidoreductase and acetyltransferase activity. Involved in patient distant metastasis-free survival in breast cancer. | None | **Yes** |
| **RASGRF2** Ras Protein Specific Guanine Nucleotide Releasing Factor 2 | Calcium-regulated nucleotide exchange factor that activates both RAS and RAS-related protein, RAC1. | Downregulated in the ovary of old mice (doi: 10.18632/aging.203150) | No |
| **GSTP1** Glutathione S-transferases | It plays an important role in detoxification by catalyzing the conjugation of many hydrophobic and electrophilic compounds with reduced glutathione.<br>Related to susceptibility to cancer. | Hypermethylated in granulosa cell tumors, causing inactivation of this suppressor gene and increasing tumor progression (doi: 10.1158/1078-0432.CCR-04-0228) | No |
| **STK17B** Serine/Threonine Kinase 17B | This nuclear protein enables ATP binding activity and protein serine/threonine kinase activity. Involved in intracellular signal transduction; positive regulation of fibroblast apoptotic process; and protein phosphorylation.<br>Highly expressed in various malignant tumors, including epithelial ovarian cancer (doi: 10.21037/atm-21-601) | None | No |
| **TBCEL** Tubulin Folding Cofactor E like | Predicted to enable alpha-tubulin binding activity, to be involved in microtubule cytoskeleton organization, post-chaperonin tubulin folding pathway and tubulin complex assembly. | Upregulate in large follicles, include in cell cycle category (doi: 10.1186/1471-2164-15-24; 10.1101/2022.10.24.513438) | No |
| **TUBA1A; TUBA3E** Tubulins (Alpha a1 and e3) | Major components of the microtubules that take part in the formation of the cytoskeleton.<br>Associated with tumor processes. | None | No |

| | | | |
|---|---|---|---|
| **HNMT** Histamine N-Methyltransferase | In charge of histamine degradation. The disregulation of its methylation involve this protein in certain cancers. | DEG in ovaries according to different environmental perturbations (doi: 10.1002/jez.b.22848) | No |
| **CADM1** Cell Adhesion Molecule 1 | Involved in cell recognition, positive regulation of cytokine production, susceptibility to NK cell mediated cytotoxicity. Implicated in cervix, prostate and ovarian tumors, where it upregulates PI3K/Akt/mTOR signaling pathway, increasing cell proliferation and migration (doi: 10.1016/j.biopha.2019.109717) | None | Yes |
| **ACP1** Acid Phospatase 1 | Hydrolyzes protein tyrosine phosphate to protein tyrosine and orthophosphate. Not related to cancer | None | No |
| **CAPG** Capping actin protein | Contributes to the control of actin-based motility in non-muscle cells. | None | No |
| **GLO1** Glyoxilase I | Linked to HLA. Participates in pyruvate metabolism and respiratory electron transport. | Involved in PCOS and decreased in old ovaries and ovulated oocytes, affecting their integrity (doi: 10.1016/j.fertnstert.2012.11.029) | No |
| **DPYD** Dihydropyrimidine Dehydrogenase | Catalyzes the reduction of uracil and thymine. Also involved in the degradation of the chemotherapeutic drug 5-fluorouracil | High expression in mural granulosa cells (doi: 10.1186/1471-2164-15-24) | No |
| **GALM** Galactose mutarotase | Cytoplasmic protein responsible for epimerization of hexose sugars such as glucose and galactose. | None | No |
| **ASMTL** Acetylserotonin O-Methyltransferase Like | Nucleoside triphosphate pyrophosphatase that hydrolyzes dTTP and UTP. May have a dual role in cell division arrest and in preventing the incorporation of modified nucleotides into cellular nucleic acids. | None | **Yes** |

**Supplementary Table 9. cMap first 100 ranked compounds that mimic the expression signature induced in KGN cells upon the elimination of FOXL2-C134W mutation.**

| Rank | Score | Name | Description |
|---|---|---|---|
| 1 | 99.93 | azathioprine | Dehydrogenase inhibitor |
| 2 | 99.86 | SIB-1757 | Glutamate receptor antagonist |
| 3 | 99.72 | **serdemetan*** | MDM inhibitor |
| 4 | 99.72 | salvinorin-a | Opioid receptor agonist |
| 5 | 99.68 | paxilline | Potassium channel blocker |
| 6 | 99.68 | NSC-119889 | Protein synthesis inhibitor |
| 7 | 99.65 | indole | aryl hydrocarbon receptor agonist |
| 8 | 99.61 | necrostatin-1 | RIPK inhibitor |
| 9 | 99.44 | PF-3845 | FAAH inhibitor |
| 10 | 99.28 | M-3M3FBS | phospholipase activator |
| 11 | 99.22 | **floxuridine*** | DNA synthesis inhibitor |
| 12 | 99.19 | HNHA | HDAC inhibitor |
| 13 | 99.19 | KU-0060648 | DNA dependent protein kinase inhibitor |
| 14 | 99.08 | APHA-compound-8 | HDAC inhibitor |
| 15 | 99.04 | benzo(a)pyrene | Carcinogen |
| 16 | 99.01 | **everolimus*** | MTOR inhibitor |
| 17 | 99.01 | **palbociclib*** | CDK inhibitor |
| 18 | 98.99 | **olaparib*** | PARP inhibitor |
| 19 | 98.98 | ISOX | HDAC inhibitor |
| 20 | 98.98 | BI-2536 | PLK inhibitor |
| 21 | 98.96 | zebularine | DNA methyltransferase inhibitor |
| 22 | 98.93 | **tretinoin*** | Retinoid receptor agonist |
| 23 | 98.91 | NCH-51 | HDAC inhibitor |
| 24 | 98.91 | bumetanide | Solute carrier family member inhibitor |
| 25 | 98.87 | givinostat | HDAC inhibitor |
| 26 | 98.87 | THM-I-94 | HDAC inhibitor |
| 27 | 98.84 | **belinostat*** | HDAC inhibitor |
| 28 | 98.84 | **vorinostat*** | HDAC inhibitor |
| 29 | 98.81 | papaverine | Phosphodiesterase inhibitor |
| 30 | 98.8 | HC-toxin | HDAC inhibitor |
| 31 | 98.8 | mycophenolic-acid | Dehydrogenase inhibitor |
| 32 | 98.78 | **panobinostat*** | HDAC inhibitor |
| 33 | 98.77 | dacinostat | HDAC inhibitor |
| 34 | 98.77 | trichostatin-a | HDAC inhibitor |
| 35 | 98.77 | apicidin | HDAC inhibitor |
| 36 | 98.77 | scriptaid | HDAC inhibitor |
| 37 | 98.77 | XMD-892 | MAP kinase inhibitor |
| 38 | 98.7 | XMD-1150 | Leucine rich repeat kinase inhibitor |
| 39 | 98.7 | cyclazosin | Adrenergic receptor antagonist |
| 40 | 98.63 | alprazolam | Benzodiazepine receptor agonist |
| 41 | 98.58 | AG-14361 | PARP inhibitor |
| 42 | 98.52 | entinostat | HDAC inhibitor |
| 43 | 98.41 | XMD-885 | Leucine rich repeat kinase inhibitor |
| 44 | 98.27 | chromomycin-a3 | DNA binding agent |
| 45 | 98.26 | methylene-blue | Guanylyl cyclase inhibitor |
| 46 | 98.24 | lofepramine | Norepinephrine reuptake inhibitor |
| 47 | 98.22 | montelukast | Leukotriene receptor antagonist |
| 48 | 98.2 | piceid | ICAM1 inhibitor |
| 49 | 98.2 | **alvocidib*** | CDK inhibitor |
| 50 | 98.17 | ER-27319 | Mediator release inhibitor |
| 51 | 98.12 | UB-165 | Acetylcholine receptor agonist |
| 52 | 98.03 | pidorubicine | Topoisomerase inhibitor |
| 53 | 98.03 | mepacrine | Cytokine production inhibitor |
| 54 | 97.99 | PHA-793887 | CDK inhibitor |
| 55 | 97.99 | hydrocotarnine | Opioid receptor antagonist |
| 56 | 97.96 | **topotecan*** | Topoisomerase inhibitor |
| 57 | 97.96 | AT-7519 | CDK inhibitor |
| 58 | 97.96 | pyroxamide | HDAC inhibitor |
| 59 | 97.93 | ketoconazole | Sterol demethylase inhibitor |
| 60 | 97.92 | cycloheximide | Protein synthesis inhibitor |
| 61 | 97.9 | lidoflazine | Calcium channel blocker |
| 62 | 97.85 | CFM-1571 | Guanylate cyclase activator |
| 63 | 97.82 | Merck60 | HDAC inhibitor |
| 64 | 97.82 | JNJ-7706621 | CDK inhibitor |
| 65 | 97.82 | **daunorubicin*** | RNA synthesis inhibitor |
| 66 | 97.78 | PIK-75 | DNA protein kinase inhibitor |
| 67 | 97.74 | **camptothecin*** | Topoisomerase inhibitor |
| 68 | 97.74 | triptolide | RNA polymerase inhibitor |
| 69 | 97.69 | prima-1-met | thioredoxin inhibitor |
| 70 | 97.67 | pirarubicin | Topoisomerase inhibitor |
| 71 | 97.64 | ZG-10 | JNK inhibitor |
| 72 | 97.64 | WT-171 | HDAC inhibitor |
| 73 | 97.6 | **mitoxantrone*** | Topoisomerase inhibitor |
| 74 | 97.57 | 5-iodotubercidin | Adenosine kinase inhibitor |
| 75 | 97.57 | KN-93 | Calcium-calmodulin dependent protein kinase inhibitor |
| 76 | 97.53 | JWE-035 | Aurora kinase inhibitor |
| 77 | 97.51 | naproxen | Cyclooxygenase inhibitor |
| 78 | 97.46 | **doxorubicin*** | Topoisomerase inhibitor |
| 79 | 97.43 | menadione | Mitochondrial DNA polymerase inhibitor |
| 80 | 97.38 | **linsitinib*** | IGF-1 inhibitor |
| 81 | 97.32 | anisomycin | DNA synthesis inhibitor |
| 82 | 97.31 | nornicotine | Acetylcholine receptor agonist |
| 83 | 97.27 | carvedilol | Adrenergic receptor antagonist |
| 84 | 97.22 | PAC-1 | Caspase activator |
| 85 | 97.21 | WAY-629 | Serotonin receptor agonist |
| 86 | 97.15 | HG-5-113-01 | Protein kinase inhibitor |
| 87 | 97.09 | AG-490 | EGFR inhibitor |
| 88 | 97.04 | clomifene | Estrogen receptor antagonist |
| 89 | 97.03 | bepridil | Calcium channel blocker |
| 90 | 96.9 | linifanib | PDGFR receptor inhibitor |
| 91 | 96.9 | PIK-90 | PI3K inhibitor |
| 92 | 96.86 | ellipticine | Topoisomerase inhibitor |
| 93 | 96.86 | JNK-9L | JNK inhibitor |
| 94 | 96.84 | SN-38 | Topoisomerase inhibitor |
| 95 | 96.83 | methyl-angolensate | Apoptosis inhibitor |
| 96 | 96.77 | PF-562271 | Focal adhesion kinase inhibitor |
| 97 | 96.73 | AZD-6482 | PI3K inhibitor |
| 98 | 96.69 | droxinostat | HDAC inhibitor |
| 99 | 96.62 | BMY-14802 | Sigma receptor antagonist |
| 100 | 96.58 | cyclopamine | Smoothened receptor antagonist |

* Drugs in common with Haltia et al. Gynecol Oncol. 2017

**Supplementary Table 10. Types of compounds that mimic the expression signature induced in KGN cells upon the elimination of FOXL2-C134W mutation.**

| Compound (cp) description | # of cp |
|---|---|
| HDAC inhibitor | 19 |
| Topoisomerase inhibitor | 8 |
| CDK inhibitor | 5 |
| Acetylcholine receptor agonist | 2 |
| Adrenergic receptor antagonist | 2 |
| Calcium channel blocker | 2 |
| Cytokine production inhibitor | 2 |
| Dehydrogenase inhibitor | 2 |
| DNA synthesis inhibitor | 2 |
| JNK inhibitor | 2 |
| Leucine rich repeat kinase inhibitor | 2 |
| Opioid receptor antagonist | 2 |
| PARP inhibitor | 2 |
| PI3K inhibitor | 2 |
| Protein synthesis inhibitor | 2 |
| Adenosine kinase inhibitor | 1 |
| Apoptosis inhibitor | 1 |
| aryl hydrocarbon receptor agonist | 1 |
| Aurora kinase inhibitor | 1 |
| Benzodiazepine receptor agonist | 1 |
| Calcium-calmodulin dependent protein kinase inhibitor | 1 |
| Carcinogen | 1 |
| Caspase activator | 1 |
| DNA binding agent | 1 |
| DNA dependent protein kinase inhibitor | 1 |
| DNA methyltransferase inhibitor | 1 |
| DNA protein kinase inhibitor | 1 |
| EGFR inhibitor | 1 |
| Estrogen receptor antagonist | 1 |
| FAAH inhibitor | 1 |

| Compound (cp) description | # of cp |
|---|---|
| Focal adhesion kinase inhibitor | 1 |
| Glutamate receptor antagonist | 1 |
| Guanylate cyclase activator | 1 |
| Guanylyl cyclase inhibitor | 1 |
| ICAM1 inhibitor | 1 |
| IGF-1 inhibitor | 1 |
| Leukotriene receptor antagonist | 1 |
| MAP kinase inhibitor | 1 |
| MDM inhibitor | 1 |
| Mediator release inhibitor | 1 |
| Mitochondrial DNA polymerase inhibitor | 1 |
| MTOR inhibitor | 1 |
| Norepinephrine reuptake inhibitor | 1 |
| PDGFR receptor inhibitor | 1 |
| Phosphodiesterase inhibitor | 1 |
| phospholipase activator | 1 |
| PLK inhibitor | 1 |
| Potassium channel blocker | 1 |
| Protein kinase inhibitor | 1 |
| Retinoid receptor agonist | 1 |
| RIPK inhibitor | 1 |
| RNA polymerase inhibitor | 1 |
| RNA synthesis inhibitor | 1 |
| Serotonin receptor agonist | 1 |
| Sigma receptor antagonist | 1 |
| Smoothened receptor antagonist | 1 |
| Solute carrier family member inhibitor | 1 |
| Sterol demethylase inhibitor | 1 |
| thioredoxin inhibitor | 1 |
